# *Arabidopsis* ORP2A positively regulates glucose signaling by interacting with AtRGS1 and promoting AtRGS1 degradation

**DOI:** 10.1101/2022.11.23.517591

**Authors:** Qian Yu, Wenjiao Zou, Kui Liu, Jialu Sun, Yanru Chao, Mengyao Sun, Qianqian Zhang, Xiaodong Wang, Xiaofei Wang, Lei Ge

## Abstract

Heterotrimeric GTP-binding proteins (G proteins) are a group of regulators essential for signal transmission into cells. AtRGS1 (Regulator of G protein Signaling 1) with intrinsic GTPase-accelerating protein (GAP) activity could suppress G protein and glucose signal transduction in *Arabidopsis*. However, how AtRGS1 activity is regulated is currently poor understood. Here we identified a knockout mutant *orp2a-1* (*oxysterol-binding protein* (*OSBP*)*-related protein 2A*) which shows phenotypes similar to *agb1-2* (*arabidopsis g-protein beta 1*). With overexpression of ORP2A, transgenic lines display short hypocotyl, hypersensitivity to sugar and lower intracellular AtRGS1 level than control. Consistently, ORP2A shows interaction with AtRGS1 *in vitro* and *vivo*. Tissue specificity of ORP2A with two alternative protein forms imply its functions in organ size and shape controlling. Bioinformatic data and phenotypes of *orp2a-1, agb1-2* and double mutant reveal genetic interactions in the regulation of G protein signaling and sugar response between ORP2A and Gβ. Both alternative splicing forms of ORP2A locate in the ER, PM (Plasma Membrane) and EPCS (ER-PM Contact Sites), and interact with VAP27-1 mediated by a FFAT-like motif *in vivo* and *vitro*. ORP2A also displays differential phosphatidyl phosphoinositide binding activity mediated by its PH domain in *vitro*. Taken together, it is suggested that *Arabidopsis* membrane protein ORP2A interacts with AtRGS1 and VAP27-1 to positively regulate G protein and sugar signaling by facilitating AtRGS1 degradation.

## INTRODUCTION

G protein signaling is a broadly conserved signal transduction mechanism in eukaryotes, which participates in almost all facets of cell biology. It has been studied relatively thorough in animals driven by medical needs (Oldham & Hamm, 2008). In the pathway the heterotrimeric GTP-binding proteins (G proteins) are critical molecular switches that transmit extracellular signals into cells facilitated by the trans-membrane G-protein-coupled receptors (GPCRs) (Gilman, 1987; Rosenbaum *et al*., 2009). G proteins have three subunits: Gα, Gβ and Gγ. When GPCR is activated by specific ligands like hormones, Gα is changed from GDP-bound to GTP-bound resulting in a conformational change. Then the Gβγ dimer is separated from Gα. Both active Gα and Gβγ dimer could mediate downstream signaling respectively. On the contrary GTPase accelerating proteins (GAPs) (such as RGS1 and phospholipase Dα1 in *Arabidopsis*, phospholipase C in mammalian cells) deactivate G proteins by accelerating the intrinsic GTPase activity of Gα and aiding GTP hydrolysis (Chen *et al*., 2003; Siderovski & Willard, 2005; Kadamur & Ross, 2013; Roy Choudhury & Pandey, 2016). Consequently, Gα would turn inactive and re-associate with Gβγ to form the heterotrimer.

Although the core components and basic properties of plant G proteins are similar to that of the metazoan system, their regulatory mechanisms seem to be rewired for plant-specific needs. Besides the common fundamental processes, G proteins in plants are also involved in diverse processes responding to plant hormones, pathogens and environmental factors, such as light, drought, and salinity (Stateczny *et al*., 2016; Pandey, 2019). Although G protein signaling is very important to growth and development, the knockout mutant of Gβ doesn’t exhibit severe developmental changes under optimal growth conditions (Lease *et al*., 2001; Ullah *et al*., 2003). The *Arabidopsis gpa1* (*g protein alpha 1*) mutants display short hypocotyls and open hooks in darkness. In light, *gpa1* shows wider leaves as a result of changes in cell proliferation pattern (Ullah *et al*., 2001). Compared to *gpa1* mutants, *agb1* (*arabidopsis g-protein beta 1*) mutants have more severely shortened hypocotyls, rounder leaves, shorter and blunter siliques (Lease *et al*., 2001; Ullah *et al*., 2003). The *agb1-2 gpa1-4* double mutant displays phenotypes similar to the single mutant *agb1-2*, which indicates that *agb1-2* is epistatic to *gpa1-4* and AGB acts downstream of GPA (Chen *et al*., 2006). Though *agg1, agg2* and *agg1 agg2* double mutant shows similar phenotypes to wild type, *agg1 agg2 agg3* triple mutant shares the morphology of *agb1* (Thung *et al*., 2012). So Gαβγ may be working together to control organ shape, and there are functional redundancy in themselves (Thung *et al*., 2012; Chakravorty *et al*., 2015).

In crops, the related G mutants exhibit similar or severer phenotypes than that in *Arabidopsis* (Fujisawa *et al*., 1999; Bommert *et al*., 2013; Urano *et al*., 2016). Studies in crops and legumes have illuminated that G protein signaling affects important agronomic traits such as plant architecture, seed size and number, and nitrogen fixation (Pandey, 2019). Recently it was found that the maize G protein β subunit regulates both meristem development and immune signaling, which can be harnessed to unleash the trade-off between growth and defense in maize (Wu *et al*., 2020). More detailed mechanistic information on the regulation of G protein signaling in plants would certainly empower future breeding applications.

The most distinctive difference between the metazoan and the plant system is that there is no canonical GPCR in plants. The plant Gα can self-activate without the help of GPCR (Johnston *et al*., 2007; Urano *et al*., 2012a). RGS (Regulator of G-protein Signaling) proteins are the key GAPs, which speed up the GTPase activity of Gα hence negatively regulates G protein signaling. AtRGS1 which has one member in *Arabidopsis* has a N-terminal seven-transmembrane (7-TM) domain that resembles GPCR and a cytosolic RGS box domain in the C-terminal (Chen *et al*., 2003). All other plant RGS proteins show high similarity to AtRGS1. It has been demonstrated that AtRGS1 protein level is sensitive to glucose, and D-glucose could accelerate the association between AtRGS1 and GPA1 (Johnston *et al*., 2007). AtRGS-related G-protein coupled glucose signal transduction pathway has been proved to be one of the three major sugar signaling pathways in plants (Jiao *et al*., 2019a). After AtRGS1 binds to GPA1, the dissociative Gβγ heterodimer recruits AtWNK8 to phosphorylate AtRGS1. Phosphorylated RGS1 undergoes endocytosis, removing the inhibition on GPA1 which activates downstream signaling (Urano *et al*., 2012b). There is a trade-off between sugar dose and treatment duration (Fu *et al*., 2014). The higher the sugar concentration, the sooner RGS1 reaches maximum internalization. Latest results proved that D-glucose induced AtRGS1 endocytosis is dependent on both Clathrin-mediated endocytosis (CME) and sterol-dependent endocytosis (SDE) (Watkins *et al*., 2021). However, the mechanism of AtRGS1 turnover is poor studied.

Oxysterol-binding protein (OSBP)-related protein (ORPs) are a type of sterol-binding proteins widely researched in yeasts and animals on a variety of cellular processes including lipid metabolism, vesicle transport and non-vesicle transport (Munro, 2003; Mesmin *et al*., 2013; Dong *et al*., 2016; Arora *et al*., 2022). ORPs generally mediate inter-organelle lipid countertransport at membrane contact sites (MCSs) (Pietrangelo & Ridgway, 2018). OSBP mediates countertransport of PI4P and cholesterol at ER-Golgi in human cells (Mesmin *et al*., 2013). Similarly, Osh6p (Oxysterol-binding homologue 6p), ORP5 and ORP8 transfer phosphatidylserine and PI4P between ER-PM in yeast and human cells (Maeda *et al*., 2013; Chung *et al*., 2015). The knockout mutant of all seven ORPs is lethal in yeast (Levine & Munro, 2001). In *Drosophila*, the loss-of-function of OSBP leads to male-sterility and defects in individualization (Ma *et al*., 2010). OSBP could repress the neurite elongation by being a target of miR-124 in Neuro-2a in mice (Gu *et al*., 2016). OSBP and ORPs are also involved in tumorigenesis, though detailed mechanism is unclear (Du *et al*., 2017).

Compared to researches in animals and yeasts, study on ORPs in plants is very limited. There are twelve members of ORPs in *Arabidopsis* (Skirpan *et al*., 2006), which are divided into four classes including class I (ORP1A, ORP1B, ORP1C, ORP1D), class II (ORP2A, ORP2B), class III (ORP3A, ORP2B, ORP3C), and class IV (ORP4A, ORP4B, ORP4C). They generally contains the ORD (OSBP-related domain) domain, the coil-coiled domain, and the signature motif of ORD domain (E(Q/K)VSHHP) plus several other different protein structures, such as the PH domain or an FFAT (two phenylalanines [FF] in an acidic tract)-like (FFATL) motif. But their functions are poorly studied. Recently, ORP2A was reported to function in autophagy regulation at ER–autophagosomal MCSs in *Arabidopsis* (Ye *et al*., 2022). In soybean, lowered expression of GmOSBP was found to result in reducing salt tolerance (Li *et al*., 2008).

Here, we identified that *Arabidopsis* lipid transport protein ORP2A interacted with AtRGS1 and VAP27-1 *in vitro* and *vivo*, functioning in AtRGS1 turnover. *Arabidopsis* ORP2A was confirmed to contain two alternative splicing forms: ORP2A with a full-length protein containing the PH domain, the Coil-coiled domain and the ORD (OSBP-related) domain, and ORP2AS (ORP2A Short) with the deletion of PH domain. Both ORP2A and ORP2AS can locate in PM, ER and EPCS. The leaves of *orp2a-1* are rounder than the wild type and the siliques are shorter and flatter, similar to G protein mutant *agb1-2*. Based on our further genetic, transcriptomic and cytological analysis, we proposed that lipid-binding protein ORP2A participates not only in organ shape controlling but also AtRGS1-related G protein and sugar signaling mediated by binding or transporting PIPs in plant.

## MATERIALS AND METHODS

### Plant materials and growth conditions

The wild-type *Arabidopsis thaliana* ecotype used in this study is Col-0. All the T-DNA insertion lines and *ORP2A* (SALK_030489c) were obtained from the Arabidopsis Biological Resource Center (ABRC, http://www.arabidopsis.org). The seeds were sterilized by bleaching (17% sodium hypochlorite, 0.04% Triton-X), then washed and spread on the medium plates with 1/2 Murashige and Skoog (MS) containing 1% (W/V) sucrose, and 0.8% (W/V) agar at pH 5.8. After synchronization for three days at 4℃, seeds are transferred to the illumination incubator and vertical cultured under the condition of 16h light, 8h dark at 22℃. The seven-day seedlings were transplanted into nutrient soil under the growth condition of16h light, 8h dark at 22℃.

### Vector construction and transformation

To test the existence of two splicing protein forms, full ORP2A genomic DNA with a 3714bp promoter was cloned and inserted into *pPZP211-3Flag. pORP2A::GFP-ORP2A* CDS was constructed to analyze subcellular localization in *Arabidopsis. pORP2A::ORP2Ag* was constructed to rescue the *orp2a-1*. To determine the expression pattern of ORP2A and ORP2AS, the 3714bp and 5936bp upstream fragments from the start cordons of ORP2A and ORP2AS were cloned into *pPZP211-GUS* vector. To generate *35S::mCherry-ORP2A, 35S:: mCherry-ORP2AS*, fragments were cloned from WT cDNA and inserted into *pPZP211-35S-mCherry* respectively. All constructs were transformed into *Agrobacterium tumefaciens* GV3101 after sequencing and then transformed into plants. All related primers used in this study are listed in Table S3.

### RNA extraction, RT-PCR and qRT-PCR

Total RNA was isolated using an Ultrapure RNA Kit (CWBIO, http://www.cwbiotech.com). Reverse transcription was performed using the RevertAid first-strand synthesis system kit (Thermo Scientific). The Quantitative PCR Q6 (ABI) was used for real-time PCR. *ACTIN2* was used as the internal reference gene. All PCR primers of this study are listed in table S3.

### GUS histochemistry

The organs of transgenic plants at different stages were incubated in GUS staining buffer (32 mM Na_2_HPO_4_, 18 mM KH_2_PO_4_, 5 mM K_3_Fe(CN)_6_, 5 mM K_4_Fe(CN)_6_, 0.5% Triton X-100, 1 mg/ml X-Gluc). The materials were transferred to be vacuumed for 15 min at 0.4 kg/cm^2^, and then transferred to 37℃ for 4 h. Non-specific staining was removed with 70% ethanol. Expression pattern was observed by Leica MZ 10F microscope.

### Confocal imaging

Two different fluorescence fusion protein constructs were transformed into *N. bentbamiana* leaf epidermal cells by *Argobacterium*-mediated transfection (Sparkes *et al*., 2006). Leaf epidermal cells were observed 2.5 days after injection. Images were captured by confocal microscope (ZEISS LSM 880). *GFP-mCherry* double-labeled materials were captured with a 488nm argon laser/BP 493-574nm filter for GFP and a 561nm argon laser/BP 578-637 nm filter for mCherry.

### Western blotting

About 200 mg tissue from five-day seedlings were used to extract protein for expression level study by 200 µL extracting buffer (50 mM Tris-HCL, 100 mM NaCl, 1%Triton-100, 10% Glycerol, 1mM PMSF, 1×protein inhibitor cocktail). Then total proteins were resolved in 8% SDS-PAGE and transferred to immobilon-P membrane (MILLIPORE cat. No. IPVH00010). GFP antibody (trans #HT801), GST antibody and Flag antibody (sigma #A8592) were used in this study.

### Yeast two hybrid assay

pGADT7 AD and pGBKT7 BD vectors were used to verify the interaction between ORP2A and VAP27-1. The plasmids were transformed into the yeast strain Y2H Gold. pPR3-SUC and pBT3-SUC vectors were used to verify the interaction between ORP2A and RGS1. The plasmids were transformed into the yeast strain NMY51. All experimental processes were operated according to protocol (Clontech).

### Pull-Down assay

GST-VAP27 was purified from E.coli BL21 by GST-beads. ORP2Ag-3Flag protein was extracted from *35S::ORP2Ag-3Flag* transformation line in extraction buffer (50 mM Tris-HCL, PH 7.5, 100 mM NaCl, 0.1% Tween-20, 10% Glycerol, 1 mM PMSF, 1×protein inhibitor cocktail). ORP2Ag-3Flag protein was incubated with GST-VAP27 in 4℃ for 2 h, after washing for three times with washing buffer (50 mM Tris-HCL, PH 7.5, 100 mM NaCl, 0.1% Tween-20, 10% Glycerol), and pull-down proteins were eluted in elution buffer (50 mM Tris-HCL, PH 7.5, 100 mM NaCl, 0.1% Tween-20, 10% Glycerol, 20 mM reduced glutathione).

### Luciferase complementation assay (LCA)

*pCAMBIA1300-nLUC* and *pCAMBIA1300-nLUC* vectors were used in this assay. All constructs were transformed into *A*.*tumefaciens* GV3101. Then two constructs were infiltrated into *N. bentbamiana* leaf epidermal cells. After 2-3 days, 0.3% D-Luciferin was infiltrated into the same region. Pictures were acquired by in vivo imaging system.

### Co-Immunoprecipitation

About 1 g tissue of 10-day old transgenic seedlings was harvested to extract total membrane proteins. The extracting process followed the manual of the Minute™ plasma membrane isolation KIT for plant (Invent. No.SM-005-P). Then the proteins were added to incubation buffer (150 mM Tris-HCl PH=7.5, 100 mM NaCL, 0.5% TritonX-100, 10% glycerol, 1 mM PMSF, 1×Protein inhibitor). About 3 μL anti-GFP antibody (Sigma-Aldrich) was added to incubation buffer and rotated at 4℃ for 2 h. 40 μL Dynabeads™ Protein G (Invitrogen) was added and rotated for another hour. Then the beads was washed three times by washing buffer (150 mM Tris-HCl PH=7.5, 100 mM NaCl, 0.1% TritonX-100, 10% glycerol, 1 mM PMSF) for ten minutes each time. At last, the beads were re-suspended in 60 μL 1×loading buffer and boiled at 100℃ for 5 min. Anti-Flag antibody (Sigma-Aldrich) and anti-GFP antibody (Transgen) were used to detect protein.

### Membrane lipid binding experiment

pGEX-4T-1 vector was used to construct *GST-ORP2A* and *GST-PH*. The proteins were purified from *E*.*coli* BL21 by GST-beads. Membrane Lipid Strips™ (product number:P-6002) and PIP Arrays™ (product number:P-6100) were purchased from Echelon Biosciences. The Protocol for PIP Strip™ membrane-type Products was followed for experimental procedure.

## RESULTS

### ORP2A functions in controlling organ development and shape formation in *Arabidopsis*

Recent evidence suggests that eukaryotic cell membranes with specific lipid composition is required for the compartmentalization of cell surface receptor signaling in immune signaling, host-pathogen interactions, cancer and cardiovascular diseases (Sezgin *et al*., 2017). And some plant cell surface receptors related to phytohormone, pathogen immunity, and heterotrimeric G-protein signaling, also needs intracellular compartmentalization to inhabit their activity (Lu *et al*., 2011; Urano *et al*., 2012; Martins *et al*., 2015; Liang *et al*., 2016).

To explore the developmental functions of membrane lipid transport proteins potentially involved in membrane protein compartmentalization in *Arabidopsis*, we phenotyped 34 T-DNA insertion lines from 8000 salk T-DNA homozygous insertion mutant collection (ABRC, http://www.arabidopsis.org), which inserted genes containing the pleckstrin homology (PH) domain, the oxysterol-binding protein (OSBP)-related protein (ORD) domain, or the phoxhomolgy (PX) domain. The *SALK_030489c* plant which had a T-DNA insertion in the fourth intron of *At4g22540(ORP2A)* (Fig. 1a), was identified as a candidate mutant with organ shape change similar to *agb1-2* (Fig.1e-f). *SALK_030489c* was referred as *orp2a-1* hereafter. The mutant *orp2a-1* as a T-DNA insertion homozygous mutant was confirmed by RT-qPCR analysis (Fig. S1a). To find out whether the T-DNA insertion disrupted the expression of *ORP2A*, RT-qPCR was performed on seven-day seedlings, and analysis showed that the expression level of *ORP2A* fragment was significantly decreased (Fig. S1b).

**Fig. 1.**
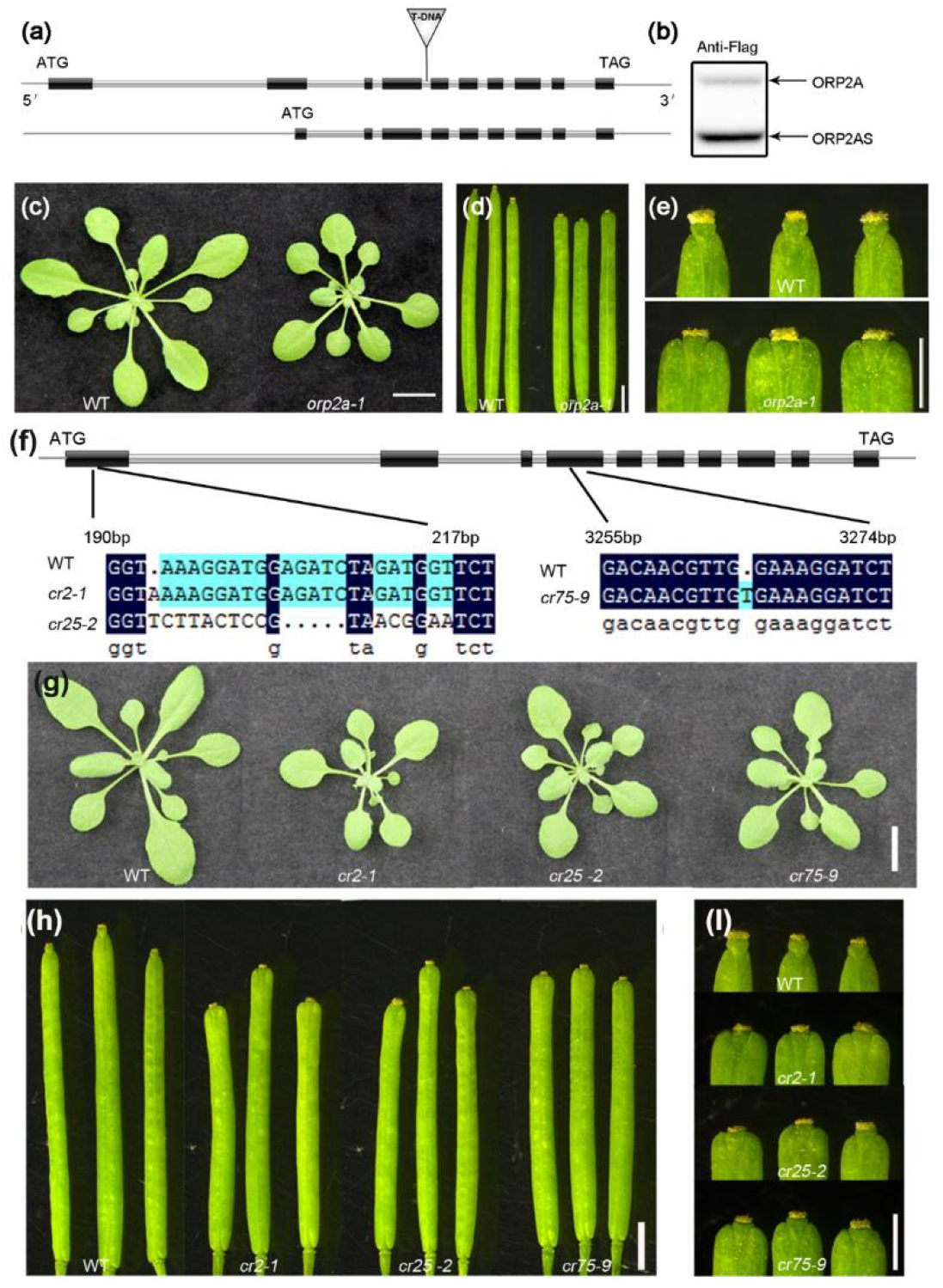
ORP2A with two splicing forms affects the morphological development of silique and leaf. (a) Schematic of full-length *ORP2A*. Inverted triangle indicates the site of T-DNA insertion in the fourth intron (the upper). Schematic of the second splicing variant *ORP2AS* (the lower). (b) Two alternative splicing forms of *ORP2A* were confirmed by Western blotting of transgenic plants with the transformation of *pORP2A:gORP2A-3*×*flag* (native promoter of *ORP2A* drives the genomic gene body of *ORP2A*). The upper band represents ORP2A-3 × Flag (Full-Length), the lower is ORP2AS-3×Flag. Antibody: anti-flag. (c) The phenotype of four-week rosette leaves of *orp2a-1* and WT. (d) The silique morphology of *orp2a-1* and WT. (e) The tip of *orp2a-1* silique (the lower) is blunter than that of WT (the upper). (f) The sequence changes of three CRISPR lines named *cr*2-1, *cr*25-1 and *cr*25-2 resulted from two different targeting locations. Changes in the left location lie in the PH domain, and the right in the ORD domain. (g) Four-week-old rosette leaves of the CRISPR lines are smaller than that of WT. (h) The silique length of three CRISPR lines are shorter than that of WT. (l)The CRISPR plant silique tips are blunter than that of WT. Bars: (c,g) 1cm; (d,h) 1mm; (e,l) 2mm.

In the website of TAIR (http://www.arabidopsis.org), it is suggest that the *ORP2A* has alternative transcriptional forms. Hence, we generated a 3×FLAG translational fusion of the *ORP2A* genomic fragment with native promoter (*pORP2A::ORP2Ag-3×Flag*) to verify the hypothesis by western blotting. The chemiluminescent assay showed two bands. This result is consistent with the prediction from TAIR website (www.arabidopsis.org) that *ORP2A* can be transcribed into two forms from alternative transcriptional start sites, resulting in two proteins with one around 100 kd (named as ORP2A), and the other about 70 kd (named as ORP2AS) (Fig. 2b).

**Fig. 2.**
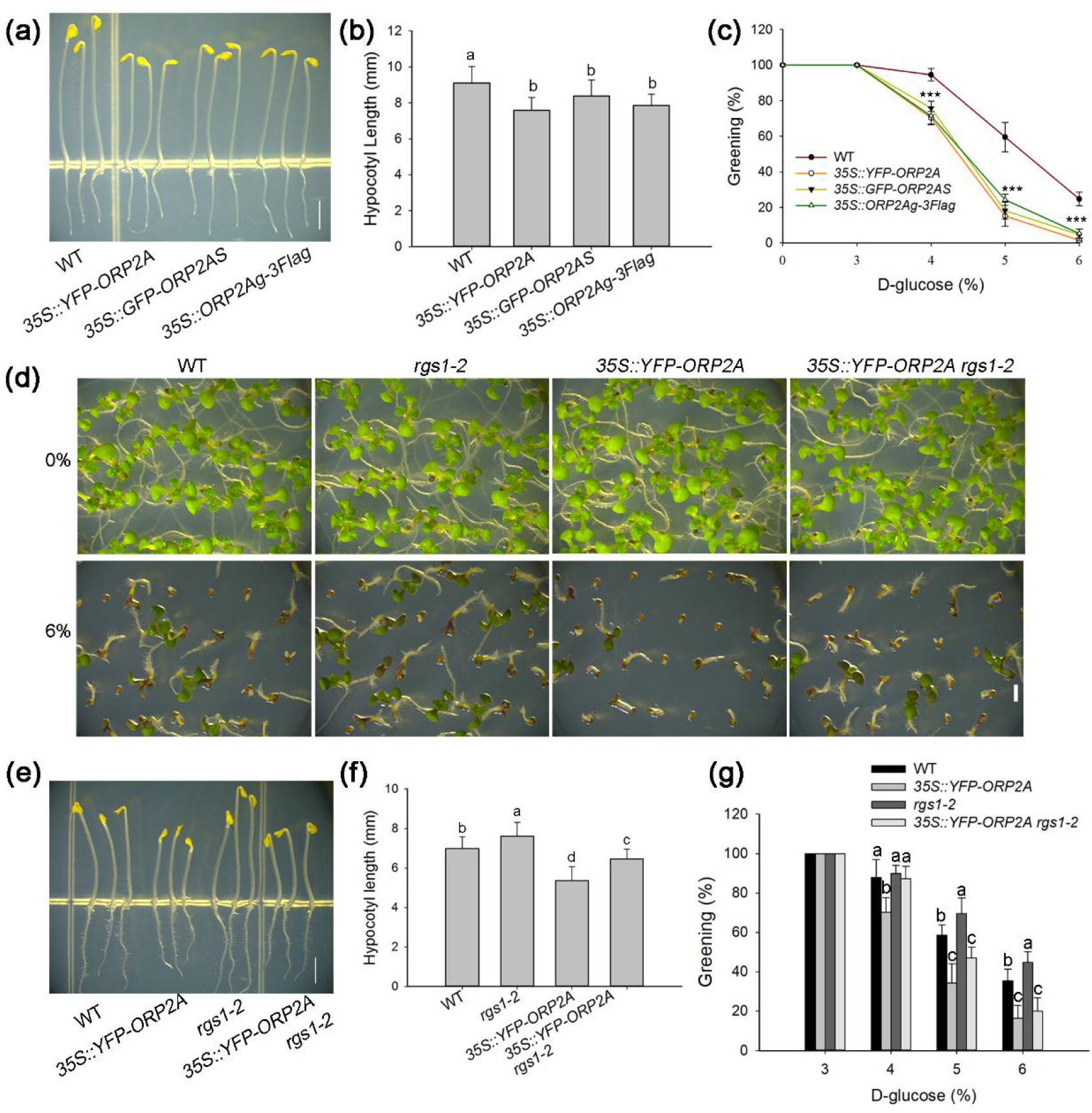
ORP2A negatively regulates hypocotyl elongation and the sugar signaling by interacting with RGS1 genetically. (a) Phenotype of hypocotyl in the WT, *35S::YFP-ORP2A, 35S::GFP-ORP2A* and *35S::ORP2Ag-3xFlag* two-days post germination in the dark. (b) The length of hypocotyl in the WT, *35S::YFP-ORP2A, 35S::GFP-ORP2A* and *35S::ORP2Ag-3xFlag* two-days post germination in the dark. (c) The greening ratio of cotyledon in the WT, *35S::YFP-ORP2A, 35S::GFP-ORP2A* and 3*5S::ORP2Ag-3xFlag* grown in medium with different D-glucose for 12 days. (d) The green seedling of WT, *35S::YFP-ORP2A, rgs1-2* and *35S::YFP-ORP2A rgs1-2* grown in medium with 0% and 6% D-glucose for 12 days. (e) The phenotype of hypocotyl in WT, *35S::YFP-ORP2A, rgs1-2* and *35S::YFP-ORP2A rgs1-2* two-days post germination in the dark. (f) The length of hypocotyl in the WT, *35S::YFP-ORP2A, rgs1-2* and *35S::YFP-ORP2A rgs1-2* two-days post germination in the dark. (g) The greening ratio of cotyledon in the WT, *35S::YFP-ORP2A, rgs1-2* and *35S::YFP-ORP2A rgs1-2* grown in medium with different D-glucose for 12 days. Bars=2mm (A, D, E). Significant differences are indicated with distinct letters in (C, E) (one-way ANOVA, Duncan’s multiple range test, p<0.05)

ORP2A is a member of Oxysterol-binding protein (OSBP)-related proteins (ORPs), which has twelve members in *Arabidopsis* (Skirpan *et al*., 2006). We rebuilt the phylogeny tree of *Arabidopsis* ORPs (Fig. S2a). Through protein sequence analysis, we found several conservative motifs similar to ORPs in animal or yeast, such as the PH domain, the ORD (OSBP-related domain) domain, the Coil-coiled domain, and the signature motif of ORD domain (E(Q/K)VSHHP) (Fig. S2a, c). ORP2A contains the PH (Pleckstrin homology) domain, the Coil-coiled domain and the ORD (OSBP-related) domain, and the PH domain was deleted in ORP2AS. In animal or yeast, the FFAT (two phenylalanines in an acidic track, EFFDAXE) motif adjacent to the coiled-coil domain is very important for its interaction with vesicle-associated membrane protein (Loewen *et al*., 2003). We did not find the same motif in all ORPs of *Arabidopsis*, but in ORP2A a similar motif (EFFNEPN) was found in a close-by position (Fig. S2b). In all the plant ORPs, no trans-membrane domain and ankyrin motif that always exist in the ORPs of animal and yeast were found (Raychaudhuri & Prinz, 2010). These data indicate that the molecular mechanism of ORPs functioning in *Arabidopsis* may be different to that in animal or yeast.

The most obvious morphology change of *orp2a-1* mutant is the rounder leaves and blunter siliques compared with WT (Fig. 1c-e). To quantify this phenotype, the fifth true leave of three-week seedlings was selected to measure length-width ratio, and the mature silique length was measured. From the results, we confirmed that the leaf of *orp2a-1* is rounder than that of WT (Fig. 1c, f), and the silique is shorter than that of WT (Fig. 2d). The tip of silique in *orp2a-1* is blunter than that of WT (Fig. 2e). The phenotypes of the mutants can be recovered by transformation of genomic DNA fragment of full-length *ORP2A* coding region with its native promoter (Fig. S3).

Since the T-DNA insertion in *orp2a-1* is in the fourth intron, it is uncertain whether it is a knock-out or knock-down mutant. Therefore, we used CRISPR-CAS9 gene-editing technology to target two different sites of ORP2A, one in the PH domain and another one in the ORD domain. We generated three CRISPR lines named *cr2-1, cr25-2* and *cr75-9* which were homozygous at the target sites. The line of *cr2-1* had a redundant adenine behind T^192^ compared to WT and composed an early stop codon. The fragment from 192-215bp in *cr*25-2 was mutated to 5’-TTCTTACTCCGTAACGGAAT-3’ to replace the fragment of 5’-TAAAGGATGGAGATCTAGATGGTT-3’of WT (Fig. 1f), while *cr75-9* line added a thymine behind C^3264^ which resulted in an early stop cordon (Fig. 1f). All the three mutants showed rounder leaves, shorter siliques and blunter silique tips similar to the phenotype of *orp2a-1* (Fig. 1g-l).

To examine when and where ORP2A and ORP2AS play their developmental roles in *Arabidopsis, pORP2A::GUS, pORP2A::GFP-ORP2A* and *pORP2AS::GUS* were constructed and transformed into wild type plants (Fig. S4-S5). The activity of *pORP2AS::GUS* is presented in the whole root-tips, primordia and leaves (Fig. S4). Because *pORP2AS::GUS* could cover the expression patterns of *pORP2A::GUS*, expression patterns of *pORP2A::GUS, pORP2A::GFP-ORP2A* were studied to explore tissue-specific activity of full-length ORP2A (Fig. S4-S5). The *pORP2A::GUS* were stained at different developmental stages. The results showed that full-length *ORP2A* was expressed in diverse tissues (Fig. S5). In young seedling root, strong GUS signal was detected in xylem, and even stronger signal was in protoxylem in both primary root tip and lateral root primordium (Fig. S5a, c, h). The confocal microscopy analysis of *pORP2A::GFP-ORP2A* also confirmed the signal in protoxylem (Fig. S5g). In seedling, *ORP2A* was primarily expressed in the margin region of leaves and cotyledons, vascular and petiole (Fig. S5a,b,e), and the strongest signal was in the growing points of leaf margin (Fig. S5d, e). Besides, *ORP2A* was selectively expressed in the ends of young silique and in filament, but not in mature silique (Fig. S5e, i). The tissue specificity of *pORP2A::GUS* indicates that the full-length ORP2A protein may participate in controlling organ shape and xylem maturation.

### ORP2A regulates hypocotyl elongation and sugar signaling by interacting with AtRGS1 genetically

Previous studies indicate that AtRGS1 has an important regulator function in G protein signaling and sugar signaling, and its mutant *rgs1-2* with defective function displayed longer hypocotyl and little response to treatment with glucose (Chen *et al*., 2003; Grigston *et al*., 2008). If ORP2A has a function in AtRGS1-associated pathway, its mutant or overexpression lines should have phenotypes related to hypocotyl elongation and sugar signaling. The hypocotyl length of *orp2a-1* in the dark and the sensitivity to D-glucose were measured, but no significant difference was detected. Then we constructed *35S::YFP-ORP2A, 35S::GFP-OPR2AS* and *35S::ORP3Ag-3Flag* overexpression transgenic lines to measure their hypocotyl length (Fig. S6). It was found that the hypocotyl length of the three independent overexpression lines were significantly shorter than that of the wild type (Fig. 2a, b). Further, we found that the overexpression lines showed a hypersensitive phenotype to glucose (Fig. 2c). Then we also introduced *rgs1-2* into *35S::YFP-ORP2A* plant by genetic hybridization to form the *35S:: YFP-ORP2A rgs1-2* double mutant for further study. Under dark condition, the hypocotyl length of *35S::YFP-ORP2A rgs1-2* was longer than that of *35S:: YFP-ORP2A* and shorter than that of *rgs1-2* (Fig. 2e, f). Greening ratios as the parameter of plant sensitivity to various concentration of D-Glucose between WT, *35S::YFP-ORP2A, rgs1-2* and *35S::YFP-ORP2A rgs1-2*, were measured and the results indicate that overexpression of *ORP2A* counteracted the defect of *rgs1-2* (Fig. 2d, g). Taken together, it is proposed that ORP2A has a genetic interaction with RGS1 to negatively regulate hypocotyl elongation and sugar signaling.

### ORP2A interacts with AGB1 genetically to control organogenesis and sugar signaling

AGB1 was reported to be involved in the control of leaf and silique morphological development (Lease *et al*., 2001). Since the phenotype of *orp2a-1* is very similar to G protein-related mutant, we hybridized *orp2a-1* with *agb1-2* and *gpa1-4 agb1-2* double mutants to obtain *agb1-2 orp2a-1* double mutant and *gpa1-4 agb1-2 orp2a-1* triple mutant. The phenotypes of *agb1-2* were severer than those of *orp2a-1* (Fig. 3). We measured the silique tip angle by the methods in previous studies (Lease *et al*., 2001). The results showed that the silique tip angle of *agb1-2 orp2a-1* was smaller than that of either *agb1-2* or *orp2a-1*, and the silique tip angle of *gpa1-4 agb1-2 orp2a-1* was similar to that of *orp2a-1* and smaller than that of *gpa1-4 agb1-2* (Fig. 3b, c). In the aspect of silique length, *agb1-2 orp2a-1* and *gpa1-4 agb1-2 orp2a-1* were shorter than *agb1-2* and *gpa1-4 agb1-2* respectively (Fig. 3d). We also measured the ratio of length/width in the fifth leaf. The leaves of *agb1-2 orp2a-1* and *gpa1-4 agb1-2 orp2a-1* were both rounder than *orp2a-1* but similar to *agb1-2* and *gpa1-4 agb1-2* (Fig. 3a, e). These data indicate that the molecular function of ORP2A in rosette leaf and silique development possibly involves G protein signaling pathway, and works in the upstream of AGB1.

**Fig. 3.**
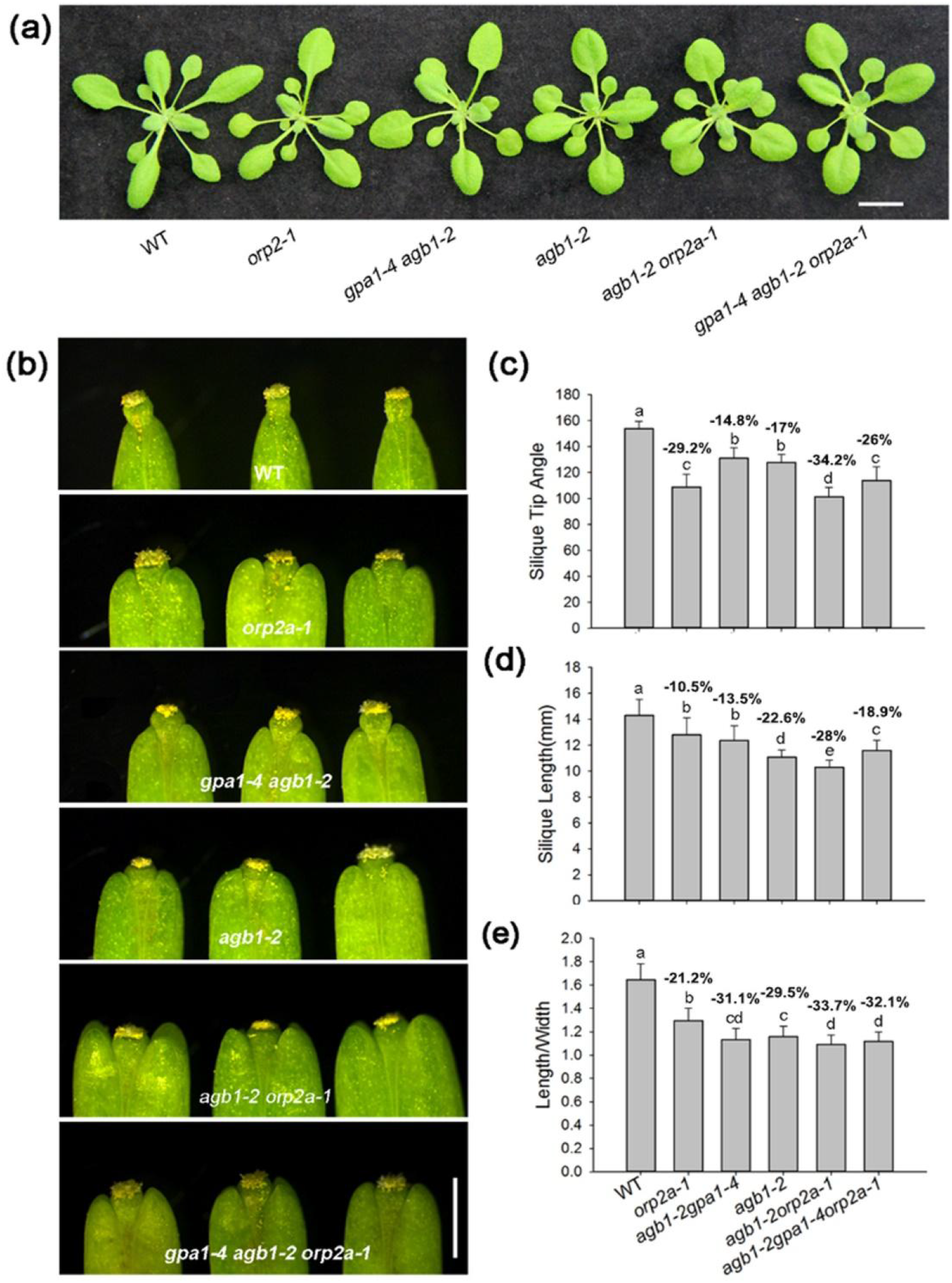
ORP2A functions in controlling organogenesis partially independent of G protein signaling. (a) Rosette leave phenotype of WT, *orp2a-1, gpa1-4 agb1-2, agb1-2, agb1-2 orp2a-1* and *gpa1-4 agb1-2 orp2a-1*. (b) The silique tip phenotype of WT, *orp2a-1, gpa1-4 agb1-2, agb1-2, agb1-2 orp2a-1* and *gpa1-4 agb1-2 orp2a-1*. (c-e) Quantitative statistics of silique tip angle (c), mature silique length (d) and length-width ratio of the fifth true leaf (e) in WT, *orp2a-1, gpa1-4 agb1-2, agb1-2, agb1-2 orp2a-1* and *gpa1-4 agb1-2 orp2a-1*. Bars, (a) 1cm; (b) 1mm. (the numbers on the column represent the percentage decline relative to WT, lower case letters indicate groups with significant difference in statistics p<0.001, Student’s *t*-test, n≥30).

To further understand the relationship between ORP2A and AGB1, we performed transcriptome sequencing on WT, *orp2a-1, agb1-2* and a*gb1-2 orp2a-1*. Through comparisons of transcript expression levels between WT and the three mutants respectively, 139 differentially expressed genes (DEGs) were found in WT vs *orp2a-1* (*p*-value<0.01 and Log2 ratio≥1), 570 DEGs were found in WT vs *agb1-2* (Log2 ratio≥1 and padj<0.05), and 744 DEGs were found in WT vs *agb1-2 orp2a-1* (Log2 ratio≥1 and padj < 0.05) (Fig. 4a). From the comparisons, nearly 40% significant changed genes (52/139) in WT vs *orp2a-1* overlaps with that in WT vs *agb1-2* (Fig. 4a). The number of up-regulated genes is much bigger than the down-regulated. 83.4%, 89.3% and 78.4% of DEGs were up-regulated in WT vs *orp2a-1*, WT vs *agb1-2*, WT vs a*gb1-2 orp2a-1*, respectively. KEGG analysis was carried out using different *p*-value thresholds (0.01 for WT vs *orp2a-1* and 0.001 for WT vs *agb1-2*) to evaluate the major pathways of DEGs. Using a *p*-value threshold of 0.005, we got 43 pathways (Table S1) and 106 pathways (Table S2) in WT vs *orp2a-1* and WT vs *agb1-2* respectively (35 pathways are shared between the two datasets). The top 20 KEGG pathways affected in *orp2a-1* (18 of them also present in that of *agb1-2*) include signal transduction, cellular response to chemical stimulus pathways, which indicates that ORP2A is involved in cellular response to environmental factors and cell signaling process (Fig. 4c and Table S1). All the results indicate that ORP2A has a function in the AGB1-associated pathway.

**Fig. 4.**
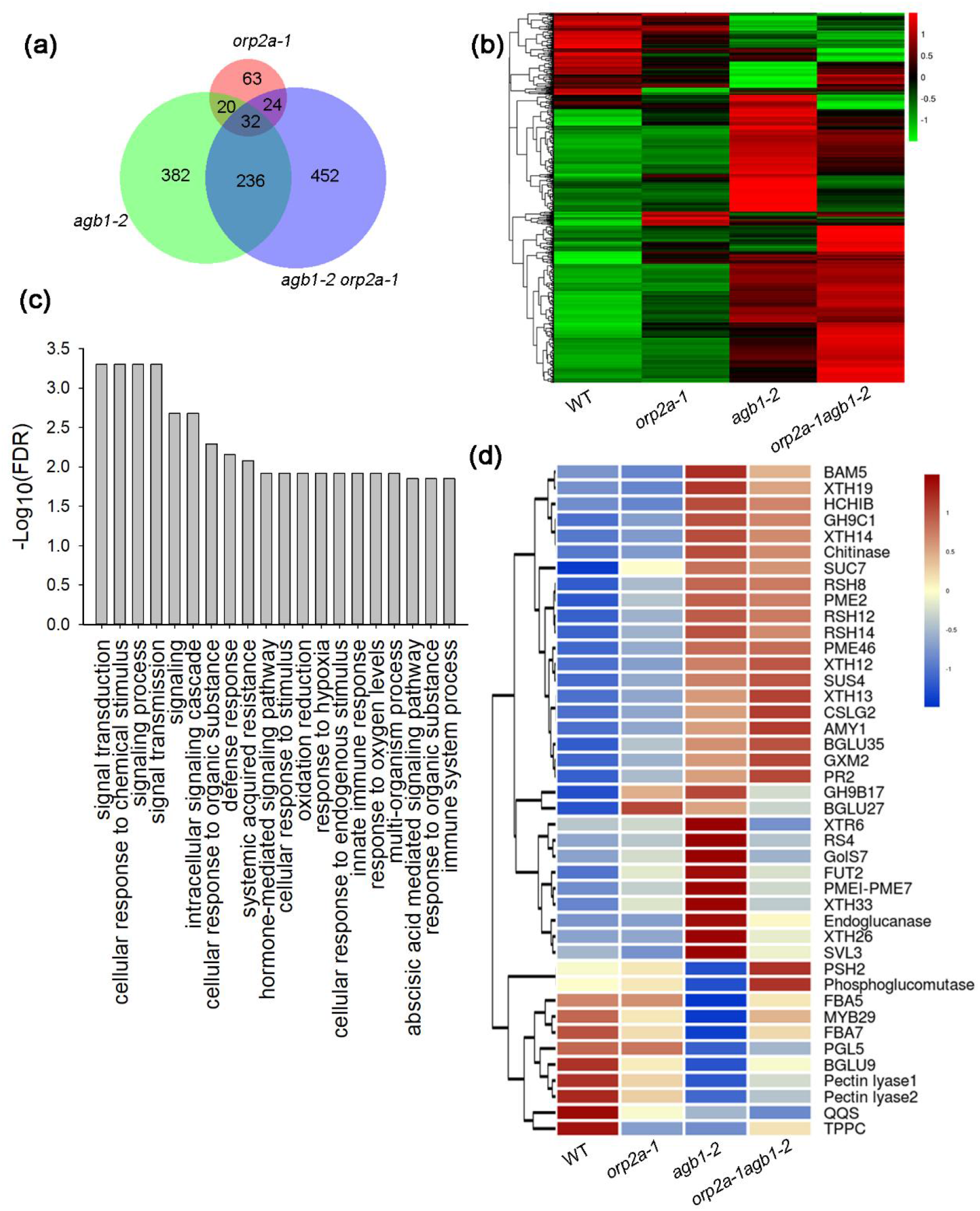
The transcriptome analysis of WT, *orp2a-1, agb1-2* and a*gb1-2orp2a-1*. (a) Venn diagram of significantly changed genes in *orp2a-1, agb1-2* and a*gb1-2orp2a-1*double mutant. (b) The heat map analysis of all significantly changed genes. (c) The top 20 KEGG of enrichment in DEGs of WT vs *orp2a-1*. (d) The heat map analysis of significantly changed genes in carbohydrate metabolic process (go:0005975).

To verify our hypothesis, the heat-map analysis of all 1453 DEGs was performed between WT and the three mutants (Fig. 4b). The map of 1453 DEGs displayed complicated pattern of transcriptional change among the four transcriptomes. Some clades of DEGs showed counteractive effects and the other ones showed synergic effects in the double mutant (Fig. 4b). But it is also found that the direction of change in most significantly changed transcripts is consistent between *agb1-2* and *orp2-1*, with a lesser extent in *orp2-1*. Because only a few genes related to sugar response are annotated in version Tair 10 (http://www.arabidopsis.org), no sugar signaling pathway was found in our analysis of DEGs. So we made the heatmap of 43 carbohydrate metabolic process genes chosen from significant DEGs in WT vs *agb1-2 (*Log2 ratio≥1 and padj < 0.05). In the heatmap, most genes showed the same tendency of changes between *agb1-2* and *orp2a-1*, but to a greater extent in WT vs *agb1-2* than that of WT vs *orp2a-1* (Fig. 4d). All those results indicate that ORP2A could work in the upstream of AGB1 in the same direction to regulate G protein signaling pathway positively, while it could also work independent of G protein pathway to regulate plant development.

### ORP2A interacts with AtRGS1 in *vitro* and *vivo*

To explore whether there is a direct interaction between ORP2A and AtRGS1, a split-ubiquitin yeast two hybrid (Y2H) for membrane protein interaction was designed. The Y2H assay verified that full-length ORP2A can interact with AtRGS1 (Fig. 5a). To test whether the two proteins interact with each other in plant cells, a Luciferase Complementation Assay (LCA) was performed in tobacco leaves. In LCA, only the PH domain of ORP2A displayed interaction with AtRGS1 (Fig. 5b). To explore whether full-length ORP2A interacts with RGS1 *in vivo*, the Co-IP assay of two proteins with the tag of GFP or 3 × Flag was examined in *Arabidopsis*. The results indicate that full-length ORP2A interacted with AtRGS1 in *vitro* and *vivo* (Fig. 5c).

**Fig.5.**
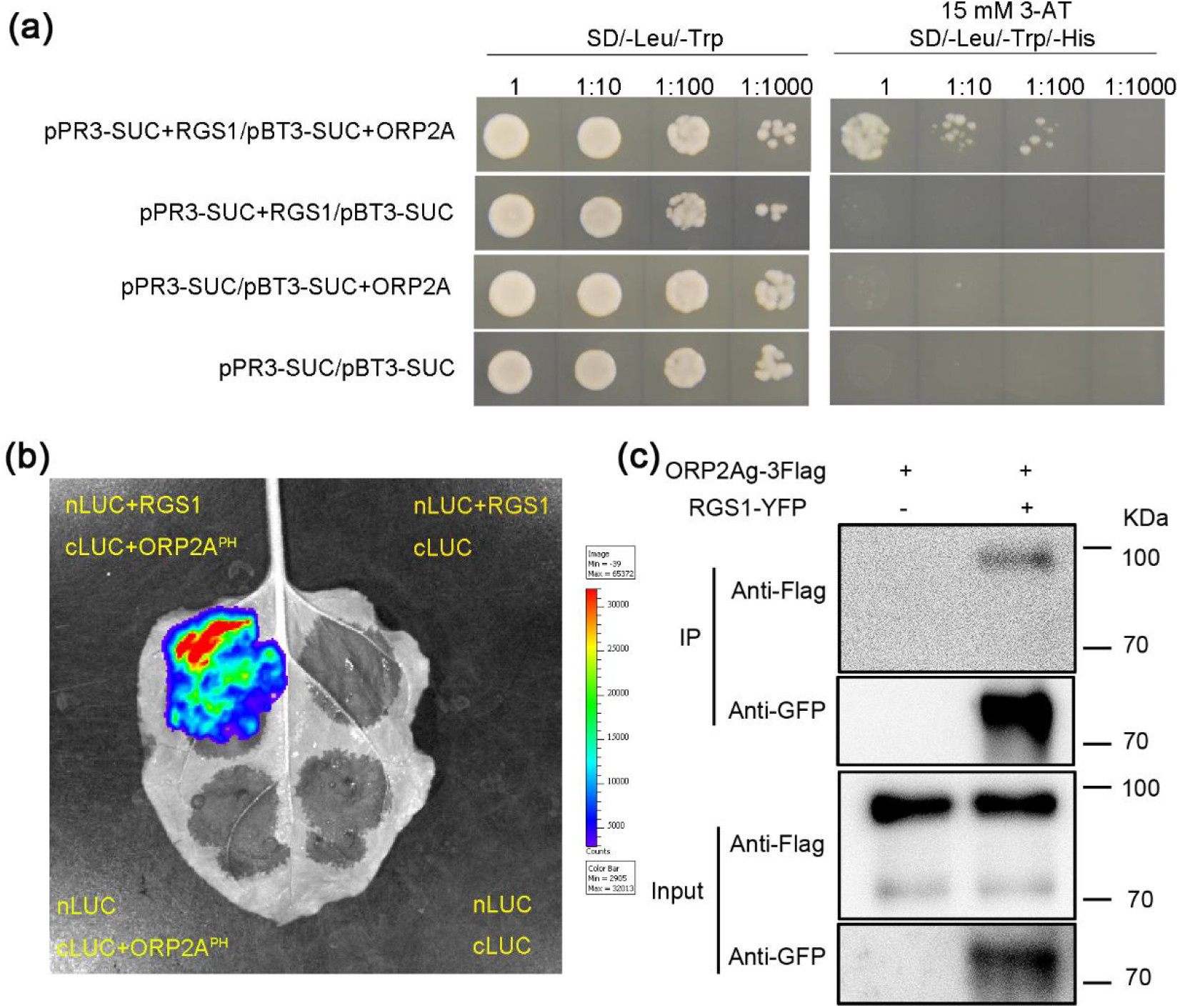
ORP2A interacts with RGS1 *in vitro* and *in vivo*. (a) ORP2A interacted with RGS1 in split-ubiquitin yeast two hybrid (Y2H). (b) The Luciferase Complementation Assay confirmed the interaction between the PH domain of ORP2A and RGS1. (c)Co-IP result of transgenic plants with the transformation of *35S:RGS1-YFP* and *35S:gORP2A-3×Flag* indicates that only full-length ORP2A interacted with RGS1 *in vitro*. RGS1-YFP was detected by anti-GFP body, and ORP2A-3Flag was detected by anti-Flag antibody.

### ORP2A localizes in PM, ER and EPCS, and interacts with VAP27-1 through FFAT-like motif

To examine the intracellular localization of the two different splicing forms of ORP2A, we cloned the full *ORP2A* CDS (722aa) and the *ORP2AS* CDS (512aa,i.e.211-722aa from full-length *ORP2A*) behind the mCherry tag driven by the 35S promoter (*35S::mCherry-ORP2A* and *35S::mCherry-ORP2AS*), and transformed them into *N. bentbamiana* leaf epidermal cells by *Argobacterium*-mediated transfection. Meanwhile, VAP27-1-GFP as the ER and EPCS marker (Wang *et al*., 2014; Wang *et al*., 2016) or PIP1;1 (PIP1;1-GFP) as the PM marker (Boursiac *et al*., 2005), were also co-expressed in the same cells by co-transfection. Both mCherry-ORP2A and mCherry-ORP2AS completely co-localized with VAP27-1-GFP in ER and EPCS, and partially co-localized with PIP1;1-GFP in PM (Fig. 6). It indicates that both forms of ORP2A localize in ER, PM and EPCS in plant cells.

**Fig. 6.**
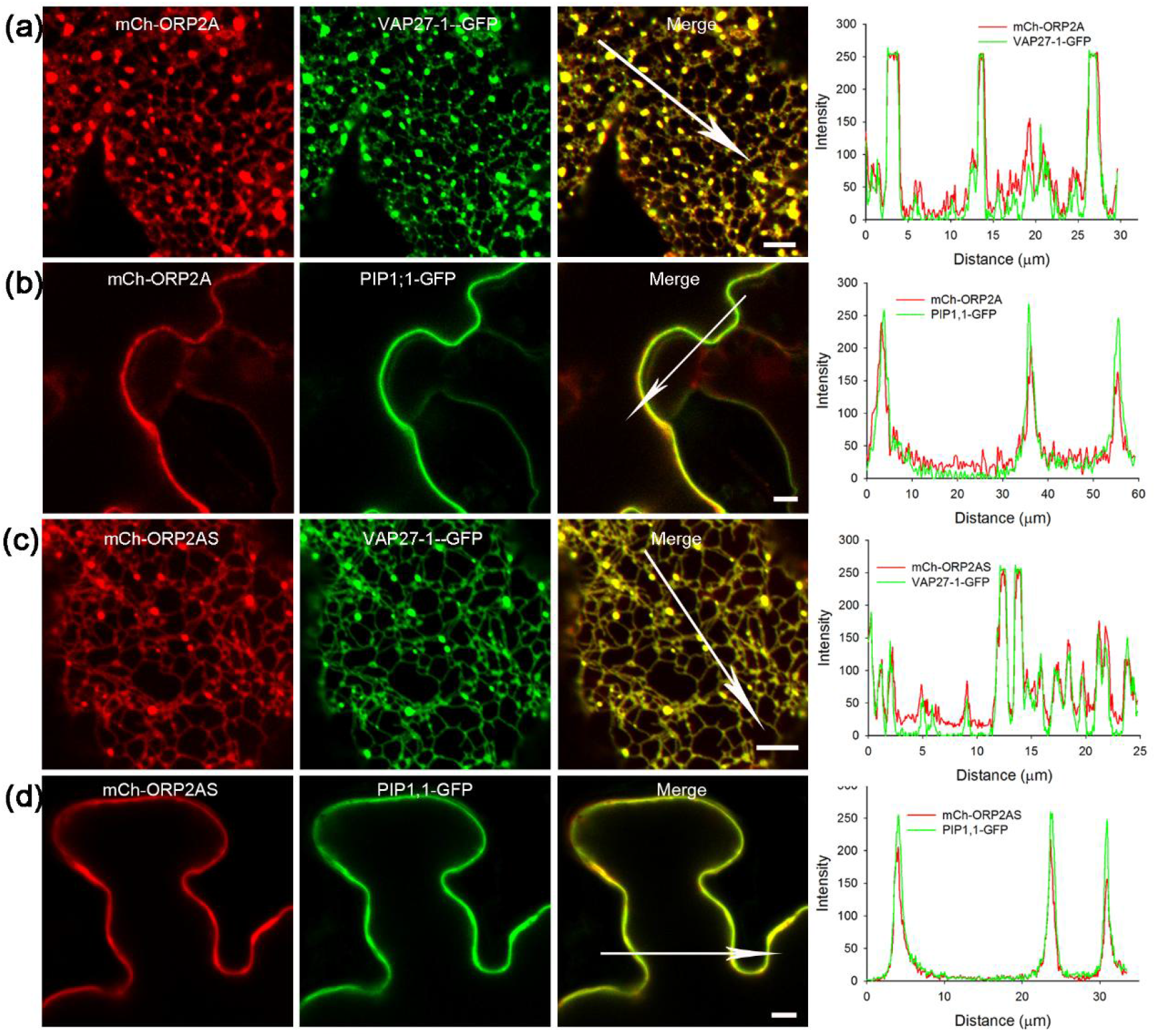
Both ORP2A and ORP2AS localize in the PM (Plasma Membrane), ER (Endoplasmic Reticulum) and PECS (PM-ER contact sites). (b-c) Confocal microscopy analyses of transiently transformed *Nicotiana benthamiana* leaf epidermal cells co-expressing mCh-ORP2A with VAP27-1-GFP (ER and PECS marker) and PIP1;1-GFP (PM marker). (b) mCh-ORP2A co-localized with VAP27-1-GFP in ER and PECS completely. (c) mCh-ORP2A partially co-localized with PIP1;1-GFP in PM. (d-e) Confocal microscopy analyses of transiently transformed *N. benthamiana* leaf epidermal cells co-expressing mCh-ORP2AS with VAP27-1-GFP and PIP1;1-GFP. (d) mCh-ORP2AS co-localized with VAP27-1-GFP in ER and PECS completely. (e) mCh-ORP2AS partially co-localized with PIP1;1-GFP in PM. Bars=10 µm.

In yeast and animal, FFAT motif is a conserved motif which targets ORPs to ER by interacting with VAPs (Loewen *et al*., 2003). Though there are not classical FFAT motifs in *Arabidopsis*, we still tested whether ORP2A could interact with VAP27-1 directly. In our analysis, full-length ORP2A did interact with VAP27-1 in the yeast two-hybrid assay (Fig. 7b). To further test which segment of ORP2A is in charge of the interaction with VAP27-1, we truncated the full ORP2A into different fragments (Fig. 7a). Two motifs containing FF amino acid were found based on the sequence analysis of ORP2A protein, with the first one (306-312aa, EFFNEPN) similar to FFAT motif of animal OSBP, while the second one (442-448aa, RFFSEKV) was located in the ORD domain. Thus, the first one (referred to as FFAT-like motif hereafter) was selected as the target motif under deletion and site mutagenesis to test for interaction between ORP2A with VAP27-1. Y2H results confirmed that ORP2A interacts with VAP27-1 through FFAT-like motif (Fig. 7b). We further confirmed that both ORP2A and ORP2AS can interact with VAP27-1 in plant cell by Luciferase Complementation Test, Pull-down and *in vivo* by Co-IP assay (Fig. 7c-e). In addition, It is found that mCherry-ORP2A^FF-AA^ with a defect FFAT-like motif could not completely co-localize with VAP27-1-GFP in EPCS (Fig. 7f). Taken the results of confocal microscopy and biochemistry together, we conclude that the FFAT-like motif is indispensable for ORP2A to interact with VAP27-1 in EPCS.

**Fig. 7.**
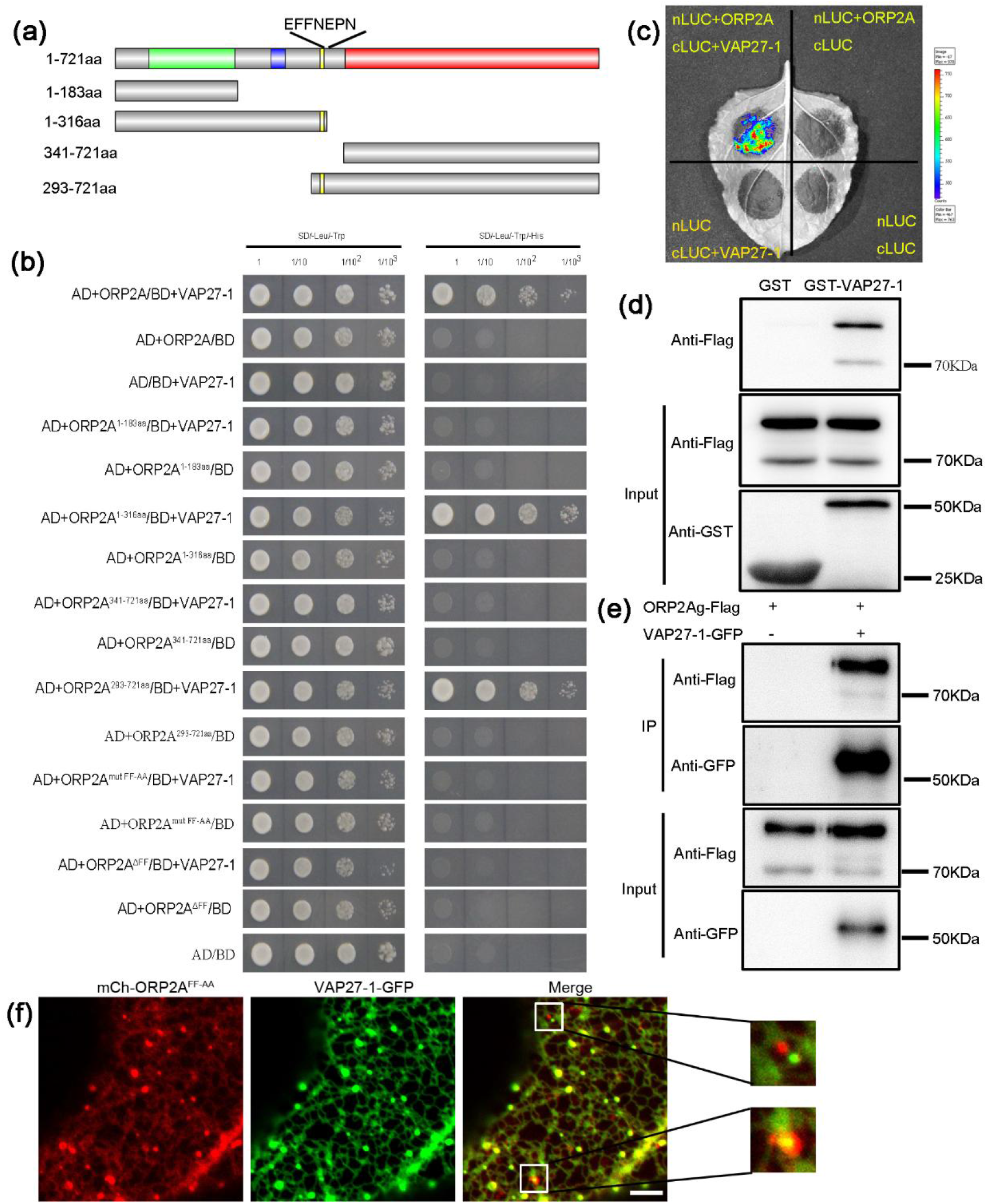
ORP2A interacted with VAP27-1 mediated by FFAT-Like motif *in vivo* and *in vitro*. (a) The schematic diagram of ORP2A protein segmentation used in the yeast two hybrid assay. Green: the PH domain, blue: the Coiled-coil domain, yellow: the FFAT-like (EFFNEPN) motif, red: the ORD domain. (b) Only those ORP2A segments containing FFAT-Like motif interacted with VAP27-1 in Y2H. (c) The LCI assay also confirmed the interaction between full-length ORP2A and VAP27-1. (d) Pull-down of ORP2A, ORP2AS with VAP27-1. The 3 × Flag-tagged of ORP2A and ORP2AS were pulled down with GST-VAP27-1 and detected by anti-Flag antibody. GST-VAP27-1 and GST were detected by anti-GST antibody. ORP2A and ORP2AS were purified from *35S:gORP2A-3 ×Flag* transformation line. (e) Co-IP result of transgenic plant with transformation of *35S:VAP27-1-GFP* and *35S:gORP2A-3×Flag* indicates that ORP2A and ORP2AS both interacted with VAP27-1 *in vitro*.VAP27-1-GFP was detected by anti-GFP body, and ORP2A-3×Flag and ORP2AS-3×Flag were detected by anti-Flag antibody. (f) Confocal images of transiently transformed *N. benthamiana* leaf epidermal cells co-expressing mCh-ORP2A^FF-AA^with VAP27-1-GFP indicate that mCh-ORP2A^FF-AA^ did not completely co-localize with VAP27-1-GFP in PECS. Bars=10 µm.

### ORP2A binds phosphatidylinositol phosphate (PIPs) differentially

Many of the PH domain in ORPs have been shown to bind phosphatidylinositol phosphate (PIPs) to a specific membrane, and the ORD domain can transfer hydroxylated cholesterol or other lipids between two membranes (Raychaudhuri & Prinz, 2010).Therefore we hypothesize that ORP2A in *Arabidopsis* may transfer some lipids between ER-PM. To verify which lipid or PIPs ORP2A binds, protein-lipid binding assays were performed on the ORP2A protein. Fig. 8a shows that ORP2A protein binds to PtdIns(4)P, PtdIns(4,5)P_2_ and PtdIns(3,4,5)P_3_, but not phosphoinositide (PI) and cholesterol. For further analysis, we purified the PH domain for assays. It was shown that the PH domain had no binding ability with PI, but had binding activities with other seven PIPs, among which, the binding ability was the strongest with PtdIns(3,5)P_2_ (Fig. 8b). Therefore, we infer that the differential binding activities of ORP2A is important for its function in PM or ER.

**Fig. 8.**
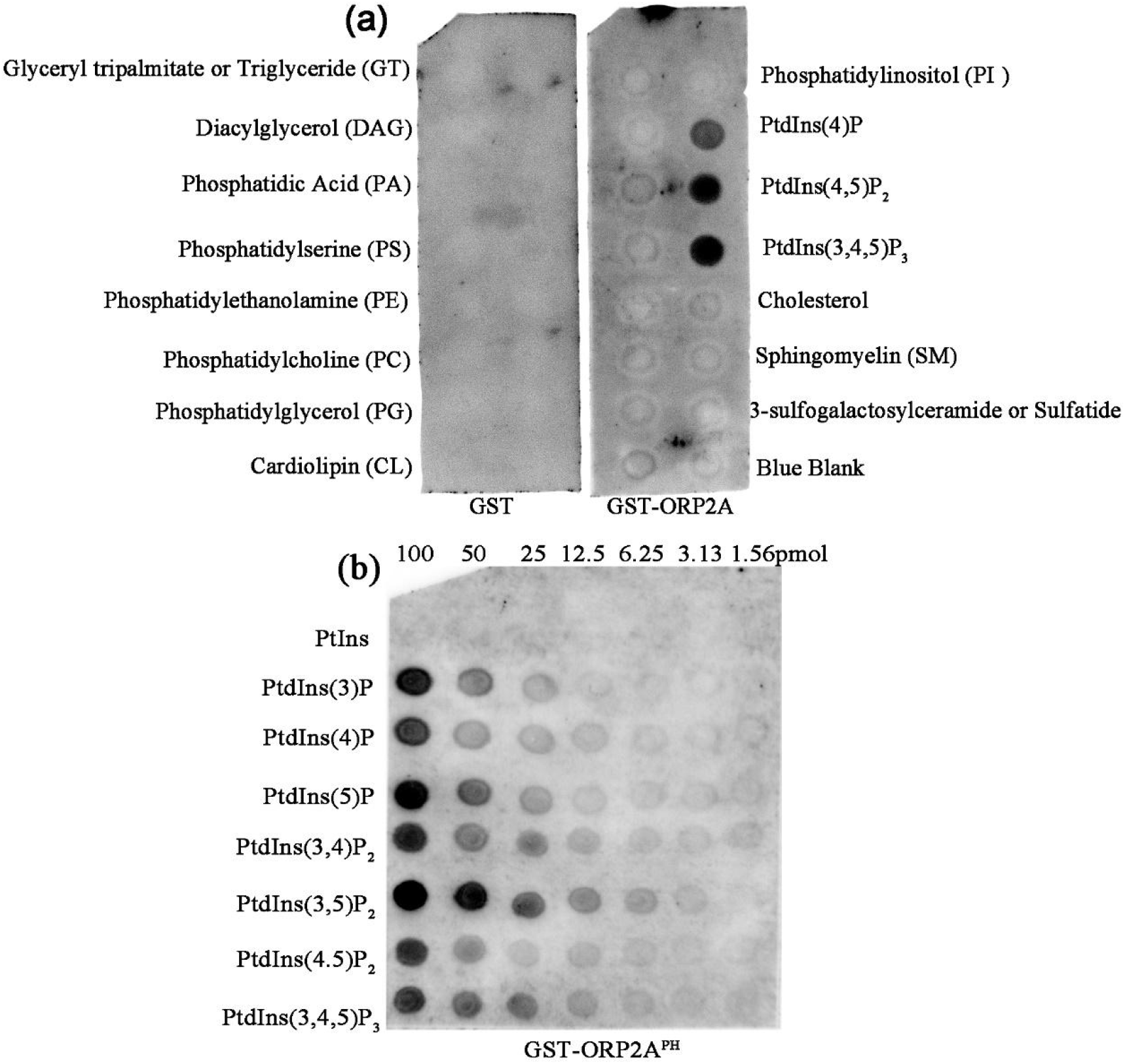
The phosphoinositide binding properties of ORP2A and ORP2A^PH^ domain. (A) The ability of the indicated GST-ORP2A fusion proteins to bind a variety of lipids analysed by a protein-lipid overlay assay. (B) The ability of the indicated GST-ORP2A^PH^ fusion proteins to bind a variety of phosphoinositides.

### ORP2A negatively regulates the intracellular AtRGS1 protein level

RGS1 could be phosphorylated by WNKs and internalized in response to D-glucose (Urano *et al*., 2012b). To verify the ORP2A function in RGS1 endocytosis or other intracellular processes, the *35S::ORP2Ag-3Flag* introduced to *35S::RGS1-YFP* transgenic lines were examined. It is observed that the fluorescence intensity in *35S::ORP2Ag-3Flag* hypocotyl was weaker than control (Fig. 9a&b). In the root-tip cells, it was found that there was no statistical difference in AtRGS1-YFP endocytosis ratio between control and *35S::ORP2Ag-3Flag* before BFA treatment for intracellular vesicle formation inhibition (Fig. 9d, e). But after BFA treatment, the YFP intensity of whole cells was decreased in *35S::ORP2Ag-3Flag* plants with any concentrations of D-glucose (Fig. 9c, d). It is also observed that the size of BFA compartments in *35S::ORP2Ag-3Flag* was smaller than that in control, and the fluorescence intensity was weaker (Fig. 9f, 9g, 9h). This result indicates that more ORP2A resulted in a faster degradation of AtRGS1-YFP after BFA treatment in the root cells. Taken the data from hypocotyl and root tip together, it is proposed that ORP2A interacted with RGS1 to promote RGS1 degradation in plant cells.

**Fig. 9.**
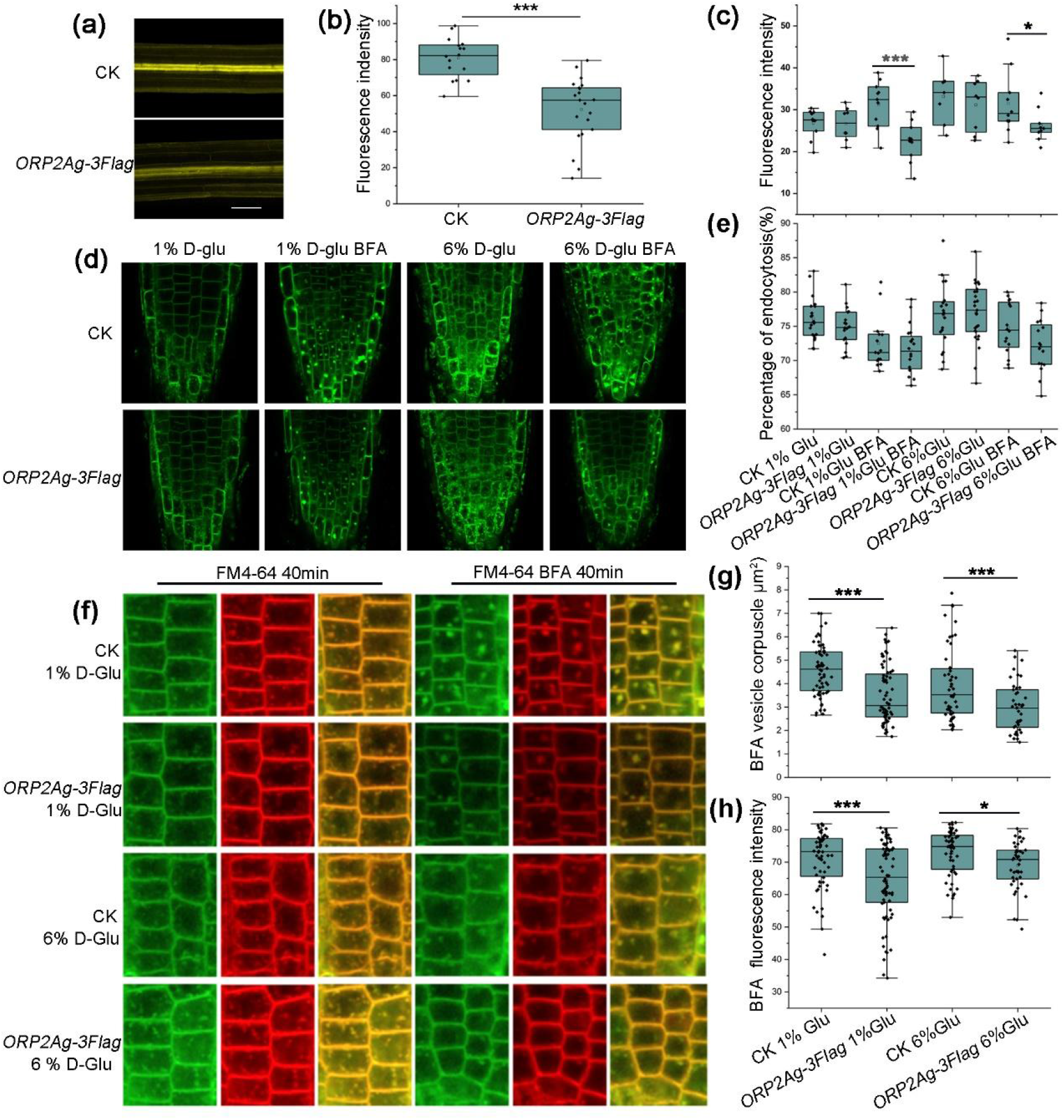
ORP2A negatively regulates the level of RGS1 protein in hypocotyle and root tip. (a) RGS1-YFP fluorescence in the hypocotyl of CK and *35S::ORP2A-3×Flag*. (b)The average fluorescence intensity of hypocotyl in (a). (c)The average fluorescence intensity of root tip in (d). (d) Confocal microscopic images of root tip cells in CK and *35S::ORP2A-3×Flag* after 40min cotreated with or without BFA and 1% or 6% D-glucose. (e) Percentage of RGS1-YFP endocytosis under the (d) condition. (f) Confocal microscopic images of root tip epidermal cells in CK and *35S::ORP2A-3×Flag* after 40min cotreated with FM4-64 with or without BFA and 1% or 6% D-glucose. (g) The area of BFA vesicle corpuscle of CK and *35S::ORP2A-3 ×Flag* after treated with 1% and 6% D-glucose. (h) The average fluorescence intensity of BFA vesicle corpuscle treated with 1% and 6% D-glucose. CK, the control plant; *ORP2A-3Flag, 35S::ORP2A-3×Flag* transgenic plant.

## DISCUSSION

### ORP2A is a molecular link between membrane lipid transport and G protein signaling

ORPs usually have more than one membrane-binding domain, such as the PH or the ORD domain, which are responsible for transferring lipid at the membrane contact sites (MCSs), and these domains are conserved from human to yeast (Raychaudhuri & Prinz, 2010). They have been confirmed to function in many cellular processes including membrane receptor signaling, vesicular trafficking, lipid metabolism, and non-vesicular sterol transfer (Raychaudhuri & Prinz, 2010). However, studies relating to ORPs in plants are very little. A recent study indicates that ORP2A has a function in autophagy regulation at ER – autophagosomal MCSs (Ye *et al*., 2022), which demonstrates its important roles in intracellular processes in plants. In our study, we found that the knockout mutant *orp2a-1* and its CRISPR lines exhibit obvious phenotypes in the change of organ shape similar to *agb1-2* (such as the rounder leaf, shorter and blunter silique) (Fig.1). ORP2A works with AGB1 and AtRGS1 in G protein and sugar signaling proved by our genetic and bioinformatical data (Fig. 2,3,4). The observed direct interaction with AtRGS1 both in *vitro* and in *vivo* provide a potential mechanism that ORP2A regulates G protein and sugar signaling in *Arabidopsis* (Fig.5).

In *Arabidopsis*, ORP2A contained both the PH, the coil-coiled, and the ORD domain (Fig. S2). We found that the PH domain of full-length ORP2A has phosphatidylinositol phosphate (PIPs) binding ability (Fig. 8b). In human and yeast, ORPs often depend on the PH domain to specifically bind certain PIPs to recruit other protein to target specific membrane (Levine & Munro, 1998; Levine & Munro, 2002). Another study in plant also proved a strong association between ORP2A and a set of PIPs (Ye *et al*., 2022). Our data further confirmed the connection between ORP2A and PIPs and pinpointed the PH domain function for ORP2A in PIP binding (Fig. 8a). GST-ORP2A did not show binding activity with other membrane lipids, in particular with cholesterol. Since we did not test with oxysterol or the other sterols, we could not exclude them from binding with ORP2A.

### Functional differences between the two alternative splicing forms of ORP2A need to be characterized

The phenomenon of alternative splicing is common among ORPs in human (Jaworski *et al*., 2001; Lehto *et al*., 2001). Human ORP1, ORP3, ORP4 and ORP9 all have been proved to have splicing variants (Lehto *et al*., 2001; Wang *et al*., 2002; Collier *et al*., 2003). For example, human ORP1 has two splicing variants, a short one that consists of the ORD domain only, and a longer one that encompassing the preceding ankyrin repeats, the PH domain and the ORD domain (Lehto *et al*., 2001; Johansson *et al*., 2003). The two variants have different intracellular localization and functions (Johansson *et al*., 2003). From our western blotting analysis, *Arabidopsis ORP2A* has two splicing variants that encode ORP2A and ORP2AS. Though the two variants share the same cellular localization, they exhibit different tissue specificity (Fig. S5&S6), and the expression level of ORP2A is lower than that of ORP2AS under native promoter (Fig. 2b). It suggests that the two splicing variants may function differently in plant development.

Our study suggests that *Arabidopsis* ORP2A’s localization in ER or PM may be mediated by binding to PIPs through the PH domain (Fig. 6 and Fig. 8). And the PH domain also mediates AtRGS1’s interaction with ORP2A (Fig.1). In animal and yeast, FFAT is a conserved motif that targets proteins to ER by interacting with VAPs (Wyles *et al*., 2002; Loewen *et al*., 2003; Lehto *et al*., 2004; Wyles & Ridgway, 2004). Although the classical FFAT motif was not found in *Arabidopsis* ORPs, both ORP2A and ORP2AS can localize in ER. We found that ORP2A and ORP2AS both interact with VAP27-1 through a FFAT-like motif, and the interaction of ORP2A and VAP27-1 is necessary for the EPCS localization of VAP27-1 (Fig. 7f), which is consistent with the findings of Jiang Lab (Ye *et al*., 2022). It suggests that the FFAT-like motif is functionally similar to FFAT. In addition, AGB1 could interacted with VAP27-1 proven by Y2H analysis (Klopffleisch *et al*., 2011). Thus, it indicates that even without the PH domain, ORP2AS may still have some function in G protein signaling, which could be independent of AtRGS1. So, the functional difference of the two alternative forms of AtORP2A needs further characterization.

### Full-length ORP2A promotes AtRGS1 degradation through autophagy

In plant, the extracellular sugar or other signals mediated by AtRGS1 need to be transmitted into cells and trigger intracellular signaling. Two working modes have been proposed to explain AtRGS1 endocytosis responsive to D-glucose, one is sterol-dependent and the other is clathrin-dependent (Watkins *et al*., 2021). However, the turnoff of AtRGS1 signal after its endocytosis is so far poor studied. It was reported that sugar starving could trigger AtRGS1 degradation mediated by autophagosome (Jiao *et al*., 2019b). PtdIns(3,5)P_2_ is proven to have crucial roles in maintaining vacuolar sorting, endocytosis of membrane proteins, autophagy, and signaling mediation in response to various cellular stresses (Shisheva, 2008). In plant, studies on the defect of PtdIns 3,5-kinase mutants of *fab1a/1b* indicate *that* PtdIns(3,5)P_2_ has functions in endomembrane homeostasis including endocytosis, vacuole formation, and vacuolar acidification (Hirano *et al*., 2011). In addition, a recent study in *Arabidopsis* shows that ORP2A mediates ER–autophagosomal MCSs and regulates autophagy through PI3P redistribution (Ye *et al*., 2022). In our study, the ratio of AtRGS1-YFP endocytosis is not significantly different between WT and *35S::ORP2Ag-3Flag* overexpression plants. But after BFA treatment, it is obvious that the size of BFA compartment in overexpression plants is smaller and YFP intensity is weaker (Fig. 9). In particular, the signal of AtRGS1-YFP in hypocotyl of overexpression plants is much weaker than the control (Fig. 9a). Given that PH domain has a higher affinity with PIPs (Fig. 8), we propose that intact ORP2A transports PIPs to AtRGS1-associated membrane lipid pool to facilitate AtRGS1 degradation through autophagy.

In summary, our study reports that ORP2A regulates G protein signaling and sugar response by interacting with RGS1 and influencing its endocytosis and degradation. Taken all data and analyses together, we propose a working model of ORP2A (Fig. S8). In the absence of ligand stimulation, RGS1 locates in the PM and inhibits G protein signaling, while ORP2A interacts with VAP27-1 located in the PM and EPCSs. When AtRGS1 turnover is needed, ORP2A would carry certain PIPs (PtdIns(3,5)P_2_ or PI3P most likely) to interact with AtRGS1, promoting AtRGS1-associated endosome sorting into vacuole. Thereby G protein signaling is activated for downstream cascade.

## ACCESSION NUMBERS

Sequence data from this work can be found in the Arabidopsis Genome Initiative or GenBank/EMBL databases under the following accession numbers: ORP2A (AT4G22540), VAP27-1 (AT3G60600), PIP1;1 (AT3G61430), AtRGS1 (AT3G26090).

## ACKNOWLEDGEMENT

We are grateful to all members of the Ge Lab for discussions on this study. This work was supported by the National Natural Science Foundation of China (grant nos 31071066, 31370325) to LG.

## AUTHOR CONTRIBUTIONS

Ge L. designed, initiated and supervised this project. Yu Q. and Zou WJ. performed most experiments. Liu K., Sun JL., Chao YR., Sun MY., Zhang QQ.,Wang XD. and Wang XF. assisted with some experiments. Zou WJ, Ge L. and Yu Q. analyzed the data and wrote the article. All authors approved the final manuscript.

## SUPPORTING INFORMATION

**Fig. S1** Identification of the recessive mutant *orp2a-1* with T-DNA insertion.

**Fig. S2** Phylogenetic analysis and domain organization of ORPs in *Arabidopsis*.

**Fig. S3** The complementation analysis of *orp2a-1* by transformation of *ORP2A* genomic gene body with native promoter.

**Fig. S4** Transcriptional activity of *ORP2A* shows high specificity in margin of young organ and protoxylem of root tip.

**Fig. S5** The comparative analysis of tissue localization of ORP2A and ORP2AS.

**Fig. S6** Identification of over-expression transgenic lines.

**Fig. S7** Working model of ORP2A function in sugar response in plant cell.

**Table S1**. KEGG pathway enrichment in WT vs *orp2a-1*.

**Table S2**. KEGG pathway enrichment in WT vs *agb1-2*.

**Table S3**. Primers used in this study.

## REFERENCES

Arora A, Taskinen JH, Olkkonen VM. 2022. Coordination of inter-organelle communication and lipid fluxes by OSBP-related proteins. Progress In Lipid Research 86: 101146.

Bommert P, Je BI, Goldshmidt A, Jackson D. 2013. The maize Galpha gene COMPACT PLANT2 functions in CLAVATA signalling to control shoot meristem size. Nature 502(7472): 555–558.

Boursiac Y, Chen S, Luu DT, Sorieul M, van den Dries N, Maurel C. 2005. Early effects of salinity on water transport in Arabidopsis roots. Molecular and cellular features of aquaporin expression. Plant Physiology 139(2): 790–805.

Chakravorty D, Gookin TE, Milner MJ, Yu Y, Assmann SM. 2015. Extra-Large G Proteins Expand the Repertoire of Subunits in Arabidopsis Heterotrimeric G Protein Signaling. Plant Physiology 169(1): 512–529.

Chen JG, Gao Y, Jones AM. 2006. Differential roles of Arabidopsis heterotrimeric G-protein subunits in modulating cell division in roots. Plant Physiology 141(3): 887–897.

Chen JG, Willard FS, Huang J, Liang J, Chasse SA, Jones AM, Siderovski DP. 2003. A seven-transmembrane RGS protein that modulates plant cell proliferation. Science 301(5640): 1728–1731.

Chung J, Torta F, Masai K, Lucast L, Czapla H, Tanner LB, Narayanaswamy P, Wenk MR, Nakatsu F, De Camilli P. 2015. INTRACELLULAR TRANSPORT. PI4P/phosphatidylserine countertransport at ORP5- and ORP8-mediated ER-plasma membrane contacts. Science 349(6246): 428–432.

Collier FM, Gregorio-King CC, Apostolopoulos J, Walder K, Kirkland MA. 2003. ORP3 splice variants and their expression in human tissues and hematopoietic cells. Dna and Cell Biology 22(1): 1–9.

Dong R, Saheki Y, Swarup S, Lucast L, Harper JW, De Camilli P. 2016. Endosome-ER Contacts Control Actin Nucleation and Retromer Function through VAP-Dependent Regulation of PI4P. Cell 166(2): 408–423.

Du X, Turner N, Yang H. 2017. The role of oxysterol-binding protein and its related proteins in cancer. Seminars In Cell & Developmental Biology.

Fu Y, Lim S, Urano D, Tunc-Ozdemir M, Phan NG, Elston TC, Jones AM. 2014. Reciprocal encoding of signal intensity and duration in a glucose-sensing circuit. Cell 156(5): 1084–1095.

Fujisawa Y, Kato T, Ohki S, Ishikawa A, Kitano H, Sasaki T, Asahi T, Iwasaki Y. 1999. Suppression of the heterotrimeric G protein causes abnormal morphology, including dwarfism, in rice. Proc Natl Acad Sci U S A 96(13): 7575–7580.

Gilman AG. 1987. G proteins: transducers of receptor-generated signals. Annual Review Of Biochemistry 56: 615–649.

Grigston JC, Osuna D, Scheible WR, Liu C, Stitt M, Jones AM. 2008. D-Glucose sensing by a plasma membrane regulator of G signaling protein, AtRGS1. Febs Letters 582(25-26): 3577–3584.

Gu X, Li A, Liu S, Lin L, Xu S, Zhang P, Li S, Li X, Tian B, Zhu X, et al. 2016. MicroRNA124 Regulated Neurite Elongation by Targeting OSBP. Molecular Neurobiology 53(9): 6388–6396.

Hirano T, Matsuzawa T, Takegawa K, Sato MH. 2011. Loss-of-function and gain-of-function mutations in FAB1A/B impair endomembrane homeostasis, conferring pleiotropic developmental abnormalities in Arabidopsis. Plant Physiology 155(2): 797–807.

Jaworski CJ, Moreira E, Li A, Lee R, Rodriguez IR. 2001. A family of 12 human genes containing oxysterol-binding domains. Genomics 78(3): 185–196.

Jiao Y, Lei W, Xu W, Chen WL. 2019a. Glucose signaling, AtRGS1 and plant autophagy. Plant Signal Behav 14(7): 1607465.

Jiao Y, Srba M, Wang J, Chen W. 2019b. Correlation of Autophagosome Formation with Degradation and Endocytosis Arabidopsis Regulator of G-Protein Signaling (RGS1) through ATG8a. Int J Mol Sci 20(17).

Johansson M, Bocher V, Lehto M, Chinetti G, Kuismanen E, Ehnholm C, Staels B, Olkkonen VM. 2003. The two variants of oxysterol binding protein-related protein-1 display different tissue expression patterns, have different intracellular localization, and are functionally distinct. Molecular Biology Of The Cell 14(3): 903–915.

Johnston CA, Taylor JP, Gao Y, Kimple AJ, Grigston JC, Chen JG, Siderovski DP, Jones AM, Willard FS. 2007. GTPase acceleration as the rate-limiting step in Arabidopsis G protein-coupled sugar signaling. Proc Natl Acad Sci U S A 104(44): 17317–17322.

Kadamur G, Ross EM. 2013. Mammalian phospholipase C. Annual Review Of Physiology 75: 127–154.

Klopffleisch K, Phan N, Augustin K, Bayne RS, Booker KS, Botella JR, Carpita NC, Carr T, Chen JG, Cooke TR, et al. 2011. Arabidopsis G-protein interactome reveals connections to cell wall carbohydrates and morphogenesis. Mol Syst Biol 7: 532.

Lease KA, Wen J, Li J, Doke JT, Liscum E, Walker JC. 2001. A mutant Arabidopsis heterotrimeric G-protein beta subunit affects leaf, flower, and fruit development. Plant Cell 13(12): 2631–2641.

Lehto M, Laitinen S, Chinetti G, Johansson M, Ehnholm C, Staels B, Ikonen E, Olkkonen VM. 2001. The OSBP-related protein family in humans. Journal Of Lipid Research 42(8): 1203–1213.

Lehto M, Tienari J, Lehtonen S, Lehtonen E, Olkkonen VM. 2004. Subfamily III of mammalian oxysterol-binding protein (OSBP) homologues: the expression and intracellular localization of ORP3, ORP6, and ORP7. Cell and Tissue Research 315(1): 39–57.

Levine TP, Munro S. 1998. The pleckstrin homology domain of oxysterol-binding protein recognises a determinant specific to Golgi membranes. Current Biology 8(13): 729–739.

Levine TP, Munro S. 2001. Dual targeting of Osh1p, a yeast homologue of oxysterol-binding protein, to both the Golgi and the nucleus-vacuole junction. Molecular Biology Of The Cell 12(6): 1633–1644.

Levine TP, Munro S. 2002. Targeting of Golgi-specific pleckstrin homology domains involves both PtdIns 4-kinase-dependent and -independent components. Current Biology 12(9): 695–704.

Li DY, Inoue H, Takahashi M, Kojima T, Shiraiwa M, Takahara H. 2008. Molecular characterization of a novel salt-inducible gene for an OSBP (oxysterol-binding protein)-homologue from soybean. Gene 407(1-2): 12–20.

Liang X, Ding P, Lian K, Wang J, Ma M, Li L, Li L, Li M, Zhang X, Chen S, et al. 2016. Arabidopsis heterotrimeric G proteins regulate immunity by directly coupling to the FLS2 receptor. Elife 5: e13568.

Loewen CJ, Roy A, Levine TP. 2003. A conserved ER targeting motif in three families of lipid binding proteins and in Opi1p binds VAP. Embo Journal 22(9): 2025–2035.

Lu D, Lin W, Gao X, Wu S, Cheng C, Avila J, Heese A, Devarenne TP, He P, Shan L. 2011. Direct ubiquitination of pattern recognition receptor FLS2 attenuates plant innate immunity. Science 332(6036): 1439–1442.

Ma Z, Liu Z, Huang X. 2010. OSBP- and FAN-mediated sterol requirement for spermatogenesis in Drosophila. Development 137(22): 3775–3784.

Maeda K, Anand K, Chiapparino A, Kumar A, Poletto M, Kaksonen M, Gavin AC. 2013. Interactome map uncovers phosphatidylserine transport by oxysterol-binding proteins. Nature 501(7466): 257–261.

Martins S, Dohmann EM, Cayrel A, Johnson A, Fischer W, Pojer F, Satiat-Jeunemaitre B, Jaillais Y, Chory J, Geldner N, et al. 2015. Internalization and vacuolar targeting of the brassinosteroid hormone receptor BRI1 are regulated by ubiquitination. Nat Commun 6: 6151.

Mesmin B, Bigay J, Moser von Filseck J, Lacas-Gervais S, Drin G, Antonny B. 2013. A four-step cycle driven by PI(4)P hydrolysis directs sterol/PI(4)P exchange by the ER-Golgi tether OSBP. Cell 155(4): 830–843.

Munro S. 2003. Cell biology: earthworms and lipid couriers. Nature 426(6968): 775–776.

Oldham WM, Hamm HE. 2008. Heterotrimeric G protein activation by G-protein-coupled receptors. Nat Rev Mol Cell Biol 9(1): 60–71.

Pandey S. 2019. Heterotrimeric G-Protein Signaling in Plants: Conserved and Novel Mechanisms. Annu Rev Plant Biol 70: 213–238.

Pietrangelo A, Ridgway ND. 2018. Bridging the molecular and biological functions of the oxysterol-binding protein family. Cellular and Molecular Life Sciences 75(17): 3079–3098.

Raychaudhuri S, Prinz WA. 2010. The diverse functions of oxysterol-binding proteins. Annu Rev Cell Dev Biol 26: 157–177.

Rosenbaum DM, Rasmussen SG, Kobilka BK. 2009. The structure and function of G-protein-coupled receptors. Nature 459(7245): 356–363.

Roy Choudhury S, Pandey S. 2016. The role of PLDalpha1 in providing specificity to signal-response coupling by heterotrimeric G-protein components in Arabidopsis. Plant Journal 86(1): 50–61.

Ruppel KM, Willison D, Kataoka H, Wang A, Zheng YW, Cornelissen I, Yin L, Xu SM, Coughlin SR. 2005. Essential role for Galpha13 in endothelial cells during embryonic development. Proc Natl Acad Sci U S A 102(23): 8281–8286.

Sezgin E, Levental I, Mayor S, Eggeling C. 2017. The mystery of membrane organization: composition, regulation and roles of lipid rafts. Nat Rev Mol Cell Biol 18(6): 361–374.

Shisheva A. 2008. PIKfyve: Partners, significance, debates and paradoxes. Cell Biology International 32(6): 591–604.

Siderovski DP, Willard FS. 2005. The GAPs, GEFs, and GDIs of heterotrimeric G-protein alpha subunits. Int J Biol Sci 1(2): 51–66.

Skirpan AL, Dowd PE, Sijacic P, Jaworski CJ, Gilroy S, Kao TH. 2006. Identification and characterization of PiORP1, a Petunia oxysterol-binding-protein related protein involved in receptor-kinase mediated signaling in pollen, and analysis of the ORP gene family in Arabidopsis. Plant Molecular Biology 61(4-5): 553–565.

Sparkes IA, Runions J, Kearns A, Hawes C. 2006. Rapid, transient expression of fluorescent fusion proteins in tobacco plants and generation of stably transformed plants. Nat Protoc 1(4): 2019–2025.

Stateczny D, Oppenheimer J, Bommert P. 2016. G protein signaling in plants: minus times minus equals plus. Current Opinion In Plant Biology 34: 127–135.

Thung L, Trusov Y, Chakravorty D, Botella JR. 2012. Ggamma1+Ggamma2+Ggamma3=Gbeta: the search for heterotrimeric G-protein gamma subunits in Arabidopsis is over. Journal Of Plant Physiology 169(5): 542–545.

Ullah H, Chen JG, Temple B, Boyes DC, Alonso JM, Davis KR, Ecker JR, Jones AM. 2003. The beta-subunit of the Arabidopsis G protein negatively regulates auxin-induced cell division and affects multiple developmental processes. Plant Cell 15(2): 393–409.

Ullah H, Chen JG, Young JC, Im KH, Sussman MR, Jones AM. 2001. Modulation of cell proliferation by heterotrimeric G protein in Arabidopsis. Science 292(5524): 2066–2069.

Urano D, Jones JC, Wang H, Matthews M, Bradford W, Bennetzen JL, Jones AM. 2012a. G protein activation without a GEF in the plant kingdom. PLoS Genet 8(6): e1002756.

Urano D, Miura K, Wu Q, Iwasaki Y, Jackson D, Jones AM. 2016. Plant Morphology of Heterotrimeric G Protein Mutants. Plant and Cell Physiology 57(3): 437–445.

Urano D, Phan N, Jones JC, Yang J, Huang J, Grigston J, Taylor JP, Jones AM. 2012b. Endocytosis of the seven-transmembrane RGS1 protein activates G-protein-coupled signalling in Arabidopsis. Nature Cell Biology 14(10): 1079–1088.

Wang C, JeBailey L, Ridgway ND. 2002. Oxysterol-binding-protein (OSBP)-related protein 4 binds 25-hydroxycholesterol and interacts with vimentin intermediate filaments. Biochemical Journal 361(Pt 3): 461–472.

Wang P, Hawkins TJ, Richardson C, Cummins I, Deeks MJ, Sparkes I, Hawes C, Hussey PJ. 2014. The plant cytoskeleton, NET3C, and VAP27 mediate the link between the plasma membrane and endoplasmic reticulum. Current Biology 24(12): 1397–1405.

Wang P, Richardson C, Hawkins TJ, Sparkes I, Hawes C, Hussey PJ. 2016. Plant VAP27 proteins: domain characterization, intracellular localization and role in plant development. New Phytologist 210(4): 1311–1326.

Watkins JM, Ross-Elliott TJ, Shan X, Lou F, Dreyer B, Tunc-Ozdemir M, Jia H, Yang J, Oliveira CC, Wu L, et al. 2021. Differential regulation of G protein signaling in Arabidopsis through two distinct pathways that internalize AtRGS1. Sci Signal 14(695).

Wu Q, Xu F, Liu L, Char SN, Ding Y, Je BI, Schmelz E, Yang B, Jackson D. 2020. The maize heterotrimeric G protein beta subunit controls shoot meristem development and immune responses. Proc Natl Acad Sci U S A 117(3): 1799–1805.

Wyles JP, McMaster CR, Ridgway ND. 2002. Vesicle-associated membrane protein-associated protein-A (VAP-A) interacts with the oxysterol-binding protein to modify export from the endoplasmic reticulum. Journal Of Biological Chemistry 277(33): 29908–29918.

Wyles JP, Ridgway ND. 2004. VAMP-associated protein-A regulates partitioning of oxysterol-binding protein-related protein-9 between the endoplasmic reticulum and Golgi apparatus. Experimental Cell Research 297(2): 533–547.

Ye H, Gao J, Liang Z, Lin Y, Yu Q, Huang S, Jiang L. 2022. Arabidopsis ORP2A mediates ER-autophagosomal membrane contact sites and regulates PI3P in plant autophagy. Proc Natl Acad Sci U S A 119(43): e2205314119.

Yu S, Yu D, Lee E, Eckhaus M, Lee R, Corria Z, Accili D, Westphal H, Weinstein LS. 1998. Variable and tissue-specific hormone resistance in heterotrimeric Gs protein alpha-subunit (Gsalpha) knockout mice is due to tissue-specific imprinting of the gsalpha gene. Proc Natl Acad Sci U S A 95(15): 8715–8720.

