## Supplementary material for "*Arabidopsis* ORP2A positively regulates glucose signaling by interacting with AtRGS1 and promoting AtRGS1 degradation": Supplimentary files

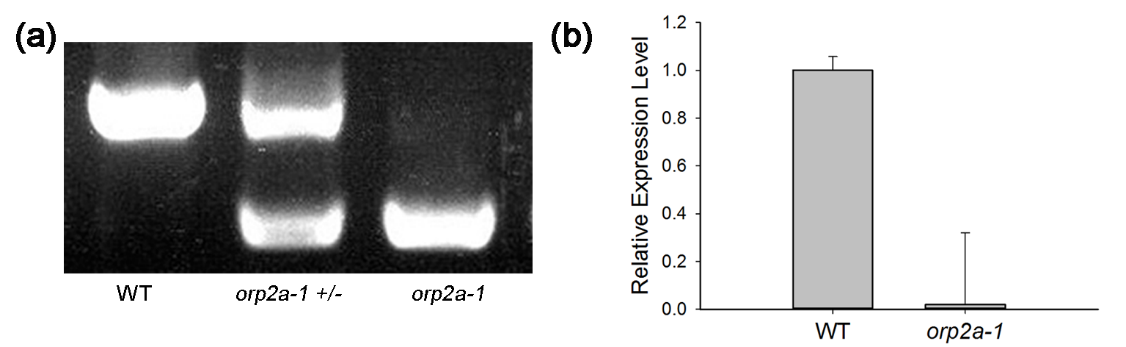

Fig. S1 Identification of the recessive mutant *orp2a-1* with a T-DNA insertion. (a) The genomic sequence of WT, heterozygous and homozygous plants were checked by PCR with the three primers (including *orp2a-1*-T-F, *orp2a-1*-T-R and LB1.3 primer on the T-DNA boundary sequence, the principle refers to http://signal.salk.edu/tdnaprimers.2.html). (b) The transcription level of ORP2A in WT and *orp2a-1* was checked by RT-qPCR. The primer sequences are showed in Table S3.

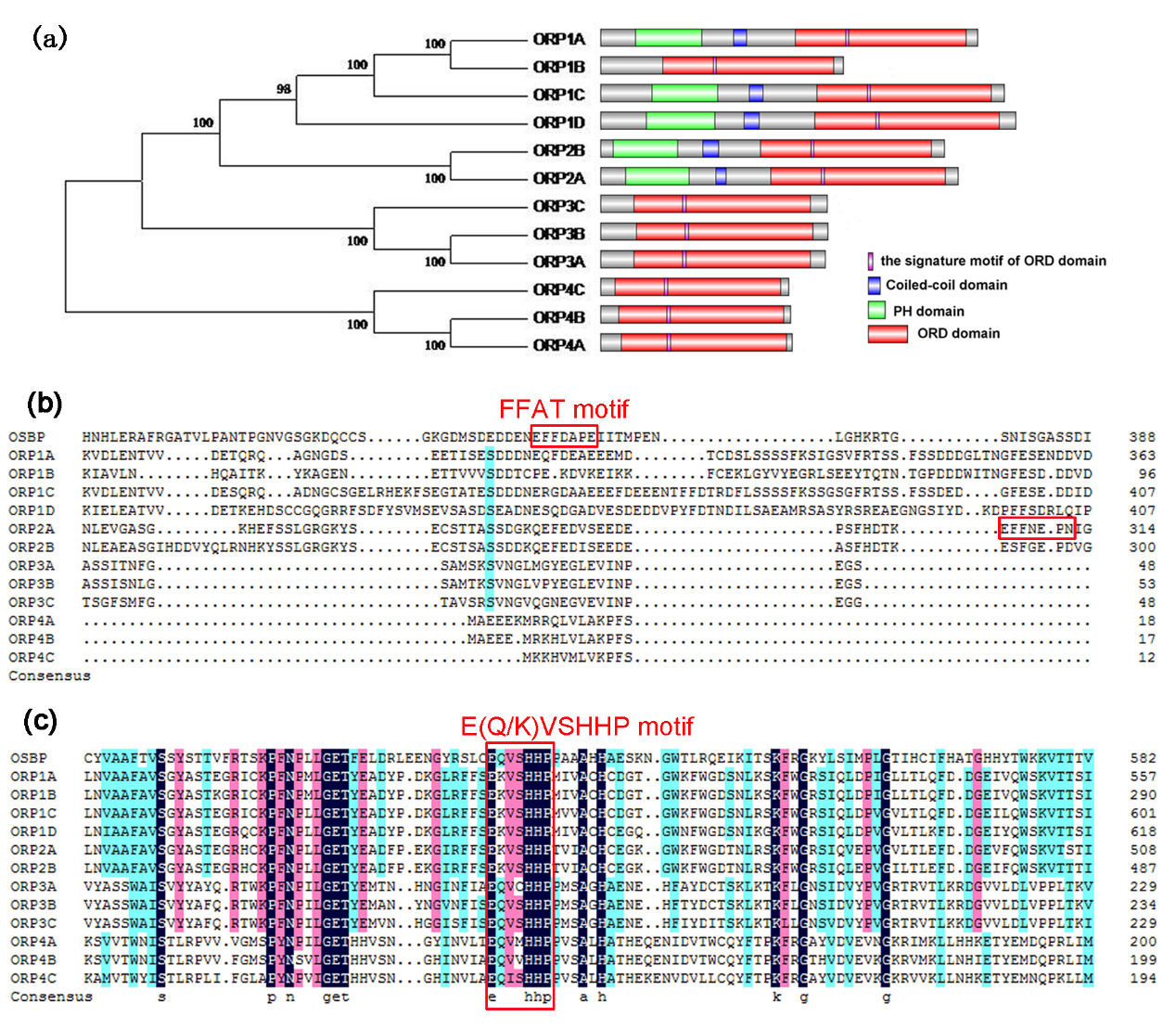

Fig. S2 Phylogenetic analysis and domain organization of ORPs in *Arabidopsis*. (a) ORPs was classified into four subfamily, classⅠ(ORP1A, ORP1B, ORP1C, ORP1D),class Ⅱ(ORP2A, ORP2B), class Ⅲ (ORP3A, ORP2B, ORP3C),class Ⅳ(ORP4A, ORP4B, ORP4C). The PH domain is shown in green, the ORD domain in red, the Coil-coiled domain in blue and the signature motif in ORD domain in purple. (b) FFAT-Like domain was found in *Arabidopsis* ORP2A. (c)The signature motif of ORD domain in ORPs is similar to that in OSBP.

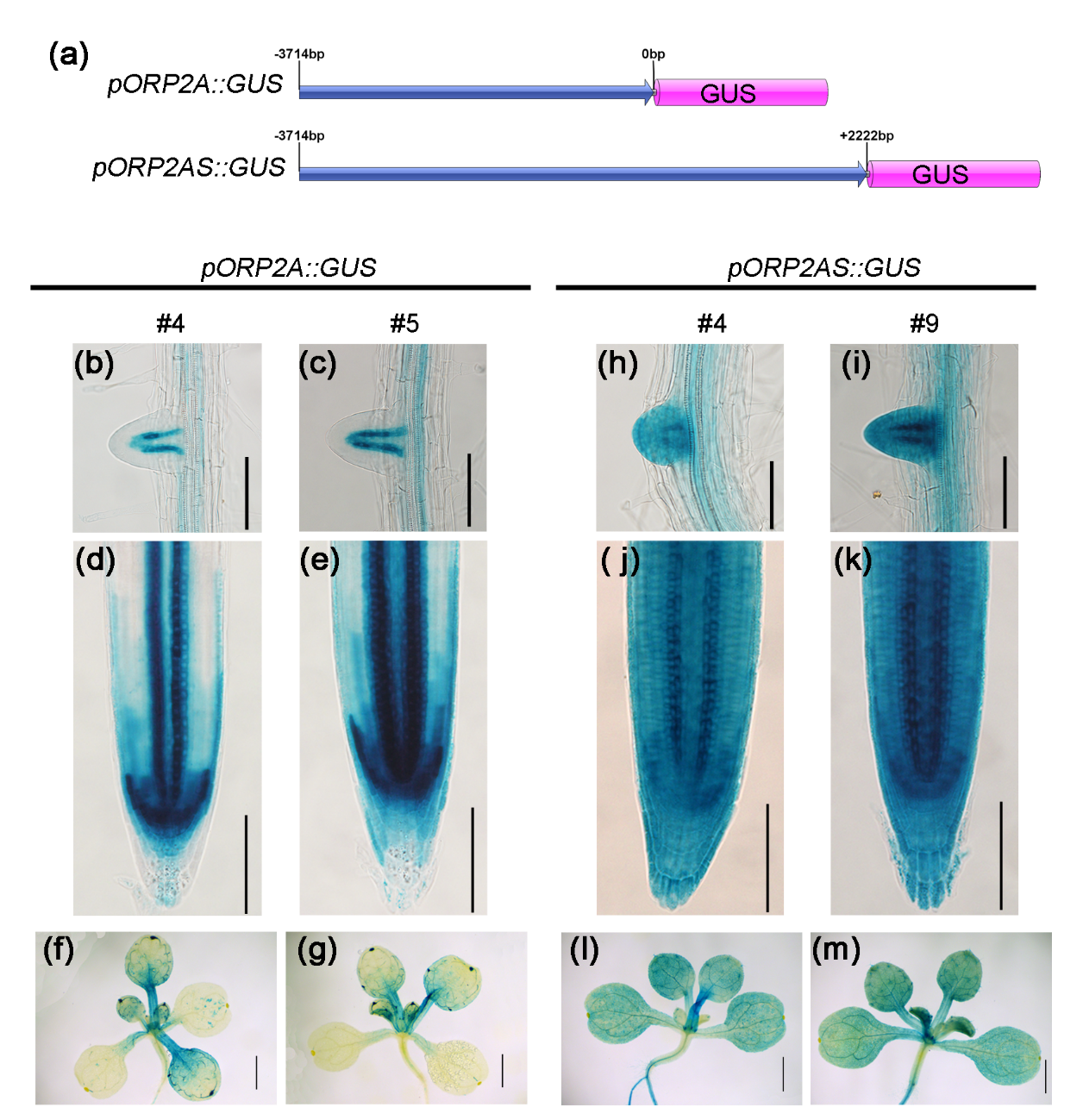

Fig. S4 The comparative analysis of tissue localization of ORP2A and ORP2AS. (a) Schematic representation of the two constructs. (b), (d) and (f) show the *pOPR2A:GUS* activity in line #4. (c), (e) and (g) are that of line #5 of *pOPR2A:GUS*. (h), (j) and (l) show the *pOPR2AS:GUS* activity in line #4 , and (i), (k) and (m) are that of line #9. Bars: (b-e, h-k) 100 µm; (f, g, l, m) 1 mm.

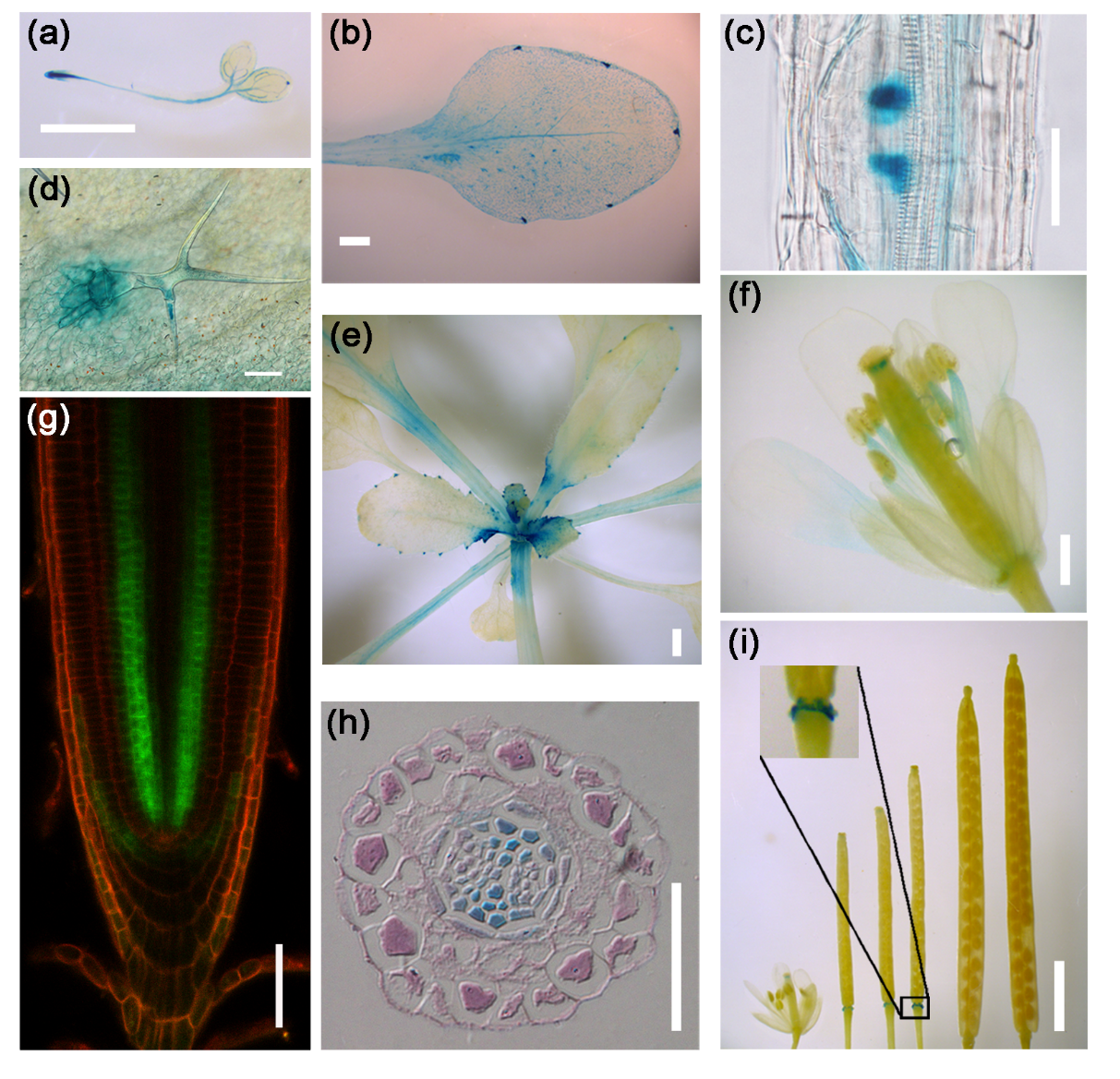

Fig. S4 Transcriptional activity of *ORP2A* shows high specificity in the margin of young organ and protoxylem of root tip. Histochemical staining of *pORP2A:GUS* plant at two-day seedlings (a), mature leaf (b), lateral root primordium (c), trichome (d), rosette leaf (e), flower (f), cross section of root tip of *pORP2A:GUS* plant (h), the different stages of silique development (i). (g) Confocal microscopy image of the root tip in *pORP2A::GFP-ORP2A* transformation line. Bars: (d, h) 100 µm; (a, b, e, i) 1 mm; (c,g) 50 µm; (f)200 µm.

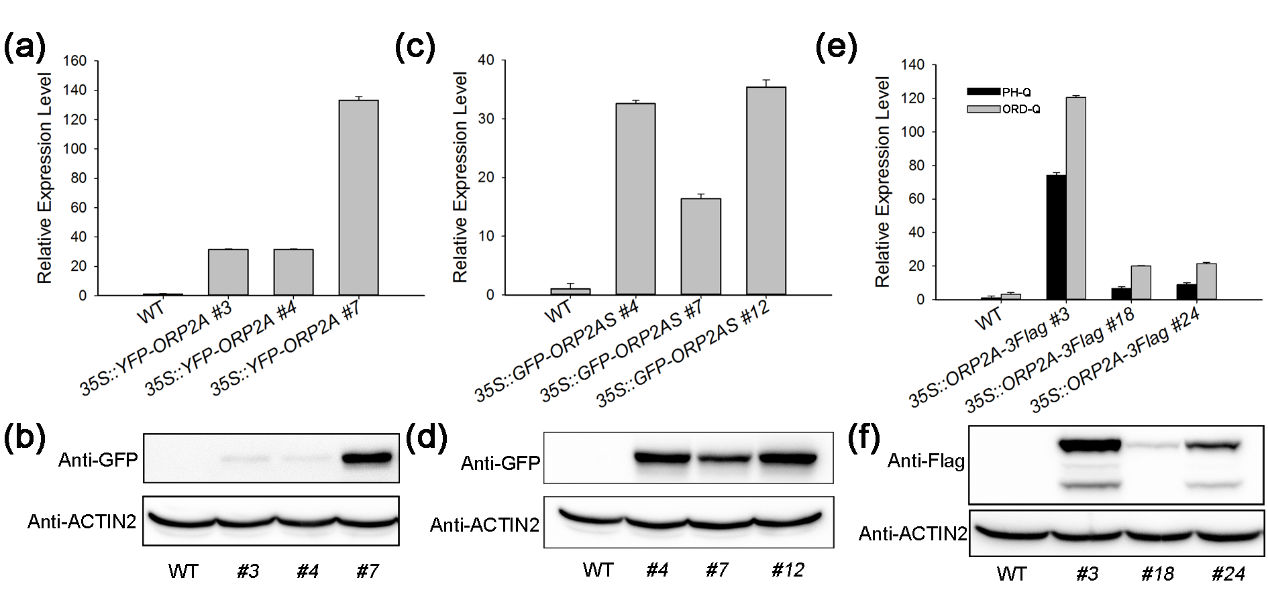

Fig. S6 Identification of over-expression transgenic lines. (a, b)The expression level of ORP2A in *35S::YFP-ORP2A* transgenic lines by RT-qPCR and western blot. (c, d) The expression level of ORP2As in *35S::GFP-ORP2AS* transgenic lines by RT-qPCR and western blot. (e, f) The relative expression level of O*RP2A* and *ORP2AS* in *35S::ORP2Ag-3Flag* transgenic lines by RT-qPCR and western blot. Identification of *35S::YFP-ORP2A* and *35S::GFP-ORP2AS* transgenic lines used the primer ORD-Q. *35S::ORP2Ag-3Flag* transgenic lines used the primers PH-Q and ORD-Q. All the primer sequences were showed in Table S3.

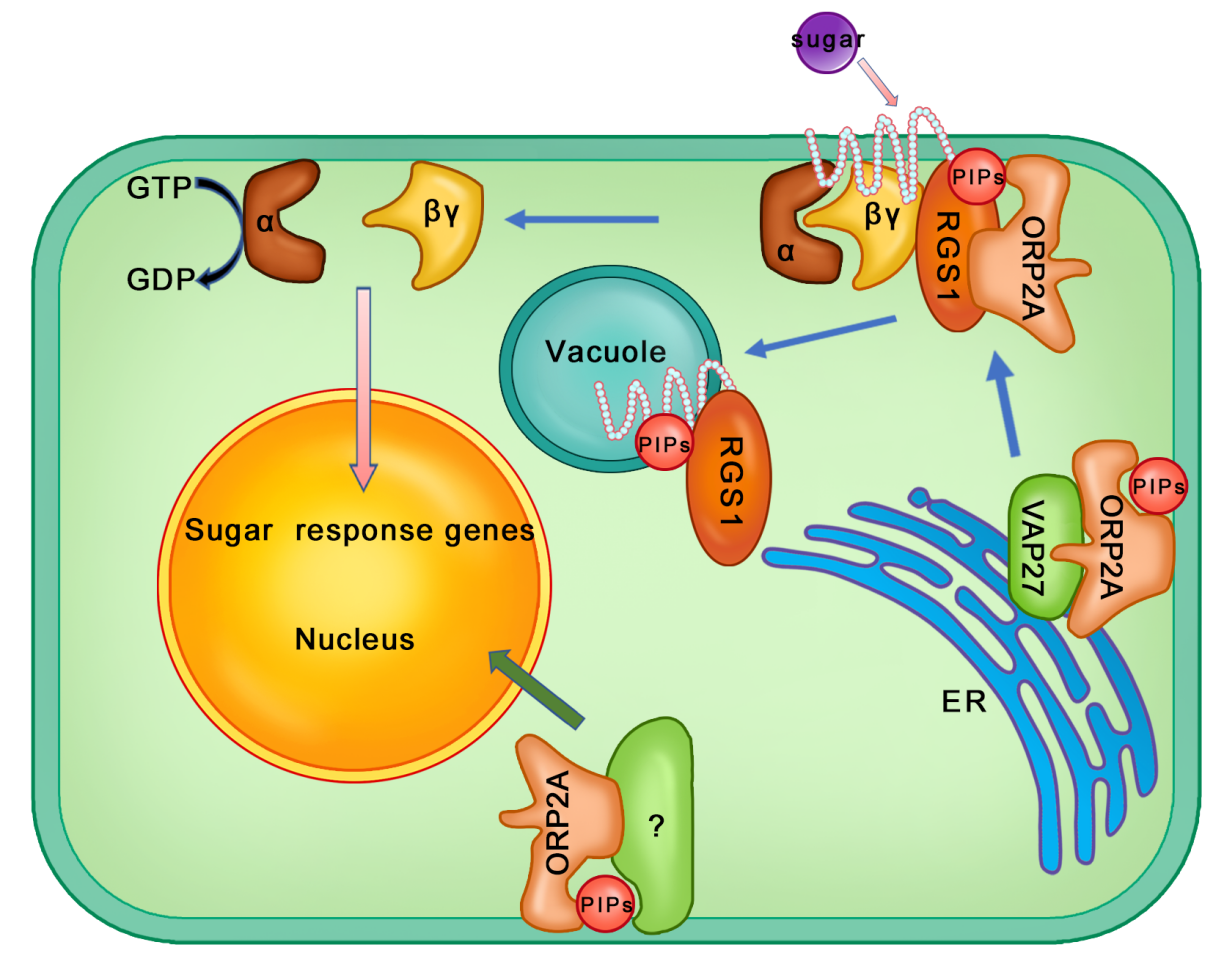

Fig. S7. The working model of ORP2A function in sugar response in plant cell*.*

| **Table S1. KEGG pathway enrichment in WT vs *orp2a-1*** | | | | | |  | |
| --- | --- | --- | --- | --- | --- | --- | --- |
| **GO term** | **Ontology** | **Description** | **Number in input list** | **Number in BG/Ref** | **p-value** | | **FDR** |
| GO:0007165 | P | signal transduction | 24 | 1670 | 2.20E-06 | | 0.0005 |
| GO:0070887 | P | cellular response to chemical stimulus | 22 | 1417 | 1.80E-06 | | 0.0005 |
| GO:0023046 | P | signaling process | 25 | 1768 | 1.80E-06 | | 0.0005 |
| GO:0023060 | P | signal transmission | 25 | 1767 | 1.70E-06 | | 0.0005 |
| GO:0023052 | P | signaling | 28 | 2376 | 1.20E-05 | | 0.0021 |
| GO:0007242 | P | intracellular signaling cascade | 19 | 1252 | 1.40E-05 | | 0.0021 |
| GO:0071310 | P | cellular response to organic substance | 18 | 1234 | 3.90E-05 | | 0.0051 |
| GO:0006952 | P | defense response | 21 | 1653 | 6.20E-05 | | 0.007 |
| GO:0009627 | P | systemic acquired resistance | 10 | 445 | 8.20E-05 | | 0.0083 |
| GO:0009755 | P | hormone-mediated signaling pathway | 11 | 600 | 0.00021 | | 0.012 |
| GO:0051716 | P | cellular response to stimulus | 25 | 2355 | 0.0002 | | 0.012 |
| GO:0055114 | P | oxidation reduction | 18 | 1364 | 0.00014 | | 0.012 |
| GO:0001666 | P | response to hypoxia | 5 | 100 | 0.00017 | | 0.012 |
| GO:0071495 | P | cellular response to endogenous stimulus | 13 | 815 | 0.00021 | | 0.012 |
| GO:0045087 | P | innate immune response | 14 | 930 | 0.00022 | | 0.012 |
| GO:0070482 | P | response to oxygen levels | 5 | 104 | 0.0002 | | 0.012 |
| GO:0051704 | P | multi-organism process | 21 | 1820 | 0.00023 | | 0.012 |
| GO:0009738 | P | abscisic acid mediated signaling pathway | 7 | 252 | 0.00029 | | 0.014 |
| GO:0010033 | P | response to organic substance | 27 | 2754 | 0.00038 | | 0.014 |
| GO:0002376 | P | immune system process | 14 | 984 | 0.00038 | | 0.014 |
| GO:0071446 | P | cellular response to salicylic acid stimulus | 8 | 351 | 0.00039 | | 0.014 |
| GO:0009863 | P | salicylic acid mediated signaling pathway | 8 | 349 | 0.00038 | | 0.014 |
| GO:0071215 | P | cellular response to abscisic acid stimulus | 7 | 267 | 0.0004 | | 0.014 |
| GO:0006955 | P | immune response | 14 | 984 | 0.00038 | | 0.014 |
| GO:0009814 | P | defense response, incompatible interaction | 10 | 536 | 0.00036 | | 0.014 |
| GO:0032870 | P | cellular response to hormone stimulus | 11 | 641 | 0.00036 | | 0.014 |
| GO:0009751 | P | response to salicylic acid stimulus | 9 | 470 | 0.00059 | | 0.02 |
| GO:0051707 | P | response to other organism | 17 | 1421 | 0.00064 | | 0.021 |
| GO:0009737 | P | response to abscisic acid stimulus | 10 | 621 | 0.0011 | | 0.034 |
| GO:0009862 | P | systemic acquired resistance, salicylic acid mediated signaling pathway | 6 | 251 | 0.0017 | | 0.05 |
| GO:0009723 | P | response to ethylene stimulus | 7 | 353 | 0.002 | | 0.058 |
| GO:0009617 | P | response to bacterium | 9 | 577 | 0.0024 | | 0.067 |
| GO:0010310 | P | regulation of hydrogen peroxide metabolic process | 5 | 187 | 0.0025 | | 0.067 |
| GO:0080010 | P | regulation of oxygen and reactive oxygen species metabolic process | 5 | 188 | 0.0026 | | 0.067 |
| GO:0009719 | P | response to endogenous stimulus | 17 | 1615 | 0.0025 | | 0.067 |
| GO:0042221 | P | response to chemical stimulus | 32 | 3978 | 0.0029 | | 0.074 |
| GO:0009725 | P | response to hormone stimulus | 15 | 1375 | 0.0033 | | 0.08 |
| GO:0019748 | P | secondary metabolic process | 14 | 1247 | 0.0035 | | 0.082 |
| GO:0050896 | P | response to stimulus | 45 | 6292 | 0.0036 | | 0.084 |
| GO:0009607 | P | response to biotic stimulus | 17 | 1687 | 0.0039 | | 0.088 |
| GO:0009697 | P | salicylic acid biosynthetic process | 5 | 209 | 0.004 | | 0.089 |
| GO:0009414 | P | response to water deprivation | 7 | 416 | 0.0048 | | 0.1 |

**Table S2. KEGG pathway enrichment in WT vs *agb1-2***

| **GO term** | **Ontology** | **Description** | **Number in input list** | **Number in BG/Ref** | **p-value** | **FDR** |
| --- | --- | --- | --- | --- | --- | --- |
| GO:0010054 | P | trichoblast differentiation | 44 | 339 | 9.30E-23 | 2.70E-19 |
| GO:0010053 | P | root epidermal cell differentiation | 44 | 350 | 2.90E-22 | 4.30E-19 |
| GO:0010015 | P | root morphogenesis | 46 | 439 | 2.60E-20 | 2.60E-17 |
| GO:0009913 | P | epidermal cell differentiation | 45 | 467 | 1.40E-18 | 7.80E-16 |
| GO:0008544 | P | epidermis development | 45 | 469 | 1.60E-18 | 7.80E-16 |
| GO:0007398 | P | ectoderm development | 45 | 469 | 1.60E-18 | 7.80E-16 |
| GO:0009814 | P | defense response, incompatible interaction | 46 | 536 | 3.50E-17 | 1.50E-14 |
| GO:0009627 | P | systemic acquired resistance | 40 | 445 | 1.10E-15 | 3.90E-13 |
| GO:0022622 | P | root system development | 46 | 595 | 1.40E-15 | 4.00E-13 |
| GO:0048364 | P | root development | 46 | 594 | 1.30E-15 | 4.00E-13 |
| GO:0010310 | P | regulation of hydrogen peroxide metabolic process | 27 | 187 | 1.90E-15 | 5.10E-13 |
| GO:0080010 | P | regulation of oxygen and reactive oxygen species metabolic process | 27 | 188 | 2.10E-15 | 5.30E-13 |
| GO:0009888 | P | tissue development | 60 | 1015 | 5.70E-15 | 1.30E-12 |
| GO:0070482 | P | response to oxygen levels | 21 | 104 | 7.90E-15 | 1.70E-12 |
| GO:0009862 | P | systemic acquired resistance, salicylic acid mediated signaling pathway | 29 | 251 | 3.20E-14 | 6.30E-12 |
| GO:0001666 | P | response to hypoxia | 20 | 100 | 4.10E-14 | 7.10E-12 |
| GO:0006952 | P | defense response | 78 | 1653 | 3.90E-14 | 7.10E-12 |
| GO:0009863 | P | salicylic acid mediated signaling pathway | 33 | 349 | 9.90E-14 | 1.60E-11 |
| GO:0071446 | P | cellular response to salicylic acid stimulus | 33 | 351 | 1.10E-13 | 1.80E-11 |
| GO:0009751 | P | response to salicylic acid stimulus | 37 | 470 | 5.50E-13 | 8.10E-11 |
| GO:0050896 | P | response to stimulus | 185 | 6292 | 1.40E-12 | 2.00E-10 |
| GO:0023052 | P | signaling | 92 | 2376 | 8.50E-12 | 1.10E-09 |
| GO:0007165 | P | signal transduction | 73 | 1670 | 9.40E-12 | 1.20E-09 |
| GO:0045087 | P | innate immune response | 51 | 930 | 1.00E-11 | 1.20E-09 |
| GO:0030154 | P | cell differentiation | 56 | 1127 | 3.50E-11 | 4.10E-09 |
| GO:0042221 | P | response to chemical stimulus | 129 | 3978 | 3.80E-11 | 4.30E-09 |
| GO:0023046 | P | signaling process | 74 | 1768 | 4.60E-11 | 4.90E-09 |
| GO:0023060 | P | signal transmission | 74 | 1767 | 4.50E-11 | 4.90E-09 |
| GO:0002376 | P | immune system process | 51 | 984 | 6.90E-11 | 6.80E-09 |
| GO:0006955 | P | immune response | 51 | 984 | 6.90E-11 | 6.80E-09 |
| GO:0051707 | P | response to other organism | 63 | 1421 | 1.70E-10 | 1.60E-08 |
| GO:0007242 | P | intracellular signaling cascade | 58 | 1252 | 2.00E-10 | 1.90E-08 |
| GO:0006800 | P | oxygen and reactive oxygen species metabolic process | 28 | 347 | 2.30E-10 | 2.10E-08 |
| GO:0042743 | P | hydrogen peroxide metabolic process | 27 | 335 | 5.00E-10 | 4.30E-08 |
| GO:0006950 | P | response to stress | 127 | 4089 | 8.60E-10 | 7.20E-08 |
| GO:0071310 | P | cellular response to organic substance | 54 | 1234 | 6.00E-09 | 4.90E-07 |
| GO:0070887 | P | cellular response to chemical stimulus | 59 | 1417 | 6.40E-09 | 5.00E-07 |
| GO:0048869 | P | cellular developmental process | 60 | 1453 | 6.30E-09 | 5.00E-07 |
| GO:0048765 | P | root hair cell differentiation | 24 | 308 | 8.50E-09 | 6.30E-07 |
| GO:0048764 | P | trichoblast maturation | 24 | 308 | 8.50E-09 | 6.30E-07 |
| GO:0048469 | P | cell maturation | 24 | 309 | 9.00E-09 | 6.50E-07 |
| GO:0051716 | P | cellular response to stimulus | 82 | 2355 | 1.60E-08 | 1.20E-06 |
| GO:0021700 | P | developmental maturation | 24 | 320 | 1.70E-08 | 1.20E-06 |
| GO:0051704 | P | multi-organism process | 68 | 1820 | 2.60E-08 | 1.70E-06 |
| GO:0009653 | P | anatomical structure morphogenesis | 67 | 1783 | 2.70E-08 | 1.80E-06 |
| GO:0048731 | P | system development | 74 | 2083 | 4.20E-08 | 2.70E-06 |
| GO:0048513 | P | organ development | 74 | 2083 | 4.20E-08 | 2.70E-06 |
| GO:0009607 | P | response to biotic stimulus | 63 | 1687 | 9.10E-08 | 5.60E-06 |
| GO:0010033 | P | response to organic substance | 89 | 2754 | 1.00E-07 | 6.10E-06 |
| GO:0065007 | P | biological regulation | 164 | 6222 | 1.60E-07 | 9.20E-06 |
| GO:0009718 | P | anthocyanin biosynthetic process | 10 | 63 | 6.70E-07 | 3.90E-05 |
| GO:0050832 | P | defense response to fungus | 21 | 342 | 2.80E-06 | 0.00016 |
| GO:0050794 | P | regulation of cellular process | 124 | 4595 | 3.20E-06 | 0.00018 |
| GO:0055114 | P | oxidation reduction | 50 | 1364 | 3.50E-06 | 0.00019 |
| GO:0009617 | P | response to bacterium | 28 | 577 | 5.10E-06 | 0.00027 |
| GO:0048468 | P | cell development | 31 | 699 | 9.00E-06 | 0.00048 |
| GO:0046283 | P | anthocyanin metabolic process | 10 | 87 | 9.30E-06 | 0.00048 |
| GO:0071554 | P | cell wall organization or biogenesis | 38 | 963 | 1.20E-05 | 0.0006 |
| GO:0009697 | P | salicylic acid biosynthetic process | 15 | 209 | 1.30E-05 | 0.00067 |
| GO:0050789 | P | regulation of biological process | 134 | 5235 | 1.60E-05 | 0.00079 |
| GO:0009719 | P | response to endogenous stimulus | 54 | 1615 | 1.80E-05 | 0.00087 |
| GO:0009696 | P | salicylic acid metabolic process | 15 | 222 | 2.60E-05 | 0.0012 |
| GO:0009620 | P | response to fungus | 24 | 499 | 2.70E-05 | 0.0013 |
| GO:0048589 | P | developmental growth | 32 | 791 | 3.70E-05 | 0.0017 |
| GO:0040007 | P | growth | 36 | 949 | 4.40E-05 | 0.002 |
| GO:0000165 | P | MAPKKK cascade | 14 | 209 | 5.20E-05 | 0.0023 |
| GO:0048856 | P | anatomical structure development | 92 | 3396 | 7.20E-05 | 0.0032 |
| GO:0009753 | P | response to jasmonic acid stimulus | 22 | 471 | 8.80E-05 | 0.0038 |
| GO:0007243 | P | protein kinase cascade | 14 | 223 | 0.0001 | 0.0043 |
| GO:0032502 | P | developmental process | 105 | 4094 | 0.00017 | 0.007 |
| GO:0042742 | P | defense response to bacterium | 19 | 394 | 0.00018 | 0.0073 |
| GO:0031323 | P | regulation of cellular metabolic process | 80 | 2928 | 0.00018 | 0.0073 |
| GO:0009595 | P | detection of biotic stimulus | 9 | 104 | 0.0002 | 0.0078 |
| GO:0019438 | P | aromatic compound biosynthetic process | 27 | 680 | 0.0002 | 0.0078 |
| GO:0031348 | P | negative regulation of defense response | 15 | 273 | 0.00023 | 0.0089 |
| GO:0009664 | P | plant-type cell wall organization | 18 | 369 | 0.00023 | 0.0089 |
| GO:0048767 | P | root hair elongation | 12 | 188 | 0.00027 | 0.01 |
| GO:0007275 | P | multicellular organismal development | 99 | 3864 | 0.00028 | 0.01 |
| GO:0009867 | P | jasmonic acid mediated signaling pathway | 15 | 282 | 0.00032 | 0.012 |
| GO:0071395 | P | cellular response to jasmonic acid stimulus | 15 | 282 | 0.00032 | 0.012 |
| GO:0006979 | P | response to oxidative stress | 23 | 582 | 0.00061 | 0.022 |
| GO:0048527 | P | lateral root development | 9 | 124 | 0.00065 | 0.023 |
| GO:0071495 | P | cellular response to endogenous stimulus | 29 | 815 | 0.00065 | 0.023 |
| GO:0009991 | P | response to extracellular stimulus | 18 | 406 | 0.00068 | 0.024 |
| GO:0009828 | P | plant-type cell wall loosening | 5 | 35 | 0.00069 | 0.024 |
| GO:0006725 | P | cellular aromatic compound metabolic process | 34 | 1022 | 0.00072 | 0.025 |
| GO:0019222 | P | regulation of metabolic process | 82 | 3186 | 0.00088 | 0.03 |
| GO:0048585 | P | negative regulation of response to stimulus | 16 | 349 | 0.00095 | 0.032 |
| GO:0032501 | P | multicellular organismal process | 99 | 4020 | 0.001 | 0.034 |
| GO:0048528 | P | post-embryonic root development | 9 | 134 | 0.0011 | 0.036 |
| GO:0071555 | P | cell wall organization | 23 | 613 | 0.0012 | 0.038 |
| GO:0031667 | P | response to nutrient levels | 16 | 367 | 0.0016 | 0.05 |
| GO:0009725 | P | response to hormone stimulus | 41 | 1375 | 0.0017 | 0.053 |
| GO:0010106 | P | cellular response to iron ion starvation | 8 | 116 | 0.0017 | 0.055 |
| GO:0051606 | P | detection of stimulus | 9 | 148 | 0.0021 | 0.065 |
| GO:0006820 | P | anion transport | 15 | 350 | 0.0025 | 0.077 |
| GO:0031668 | P | cellular response to extracellular stimulus | 16 | 388 | 0.0026 | 0.081 |
| GO:0009723 | P | response to ethylene stimulus | 15 | 353 | 0.0027 | 0.081 |
| GO:0071496 | P | cellular response to external stimulus | 16 | 389 | 0.0027 | 0.081 |
| GO:0044036 | P | cell wall macromolecule metabolic process | 14 | 319 | 0.0028 | 0.083 |
| GO:0009812 | P | flavonoid metabolic process | 12 | 251 | 0.0029 | 0.083 |
| GO:0071669 | P | plant-type cell wall organization or biogenesis | 18 | 473 | 0.0034 | 0.098 |
| GO:0048588 | P | developmental cell growth | 17 | 440 | 0.0037 | 0.11 |
| GO:0015980 | P | energy derivation by oxidation of organic compounds | 9 | 170 | 0.005 | 0.14 |

**Table S3. Primers used in this study**

| Primer’s Name | Sequence (5’-3’) | Restriction Enzyme | Vector | Purpose |
| --- | --- | --- | --- | --- |
| ORP2A-P-F | ATGAATTCCGCATATATGCAACCCAGAC | EcoRⅠ  KpnⅠ | pPZP211-GUS | pORP2A-GUS |
| ORP2A-P-R | ATGGTACCATGCAGATTTGGGGGATG |  |  |  |
| ORP2AS-P-F | ATGGTACCTCTGGATCTTACGAATGTTGC | KpnⅠ | pPZP211-pORP2A-GUS | pORP2AS -GUS |
| ORP2AS-P-R | ATGGATCCACCCTCTTCGTGCAACCG | BamHⅠ |  |  |
| ORP2A-GK-F | ATGGTACCCTCTGGATCTTACGAATGTTGC | KpnⅠ | pPZP211-35S-3Flag | 35S::ORP2Ag-3Flag |
| ORP2A-GB-R | ATGGATCCAGCGGCGTCCGCTAGCTCTT | BamHⅠ |  |  |
| ORP2A-CS-F | ATGTCGACATGCGGGTTAAAGAGTTAC | SalⅠ | pPZP211-35S-mCherry | 35S:mCherry-ORP2A |
| ORP2A-CP-R | ATCTGCAGCTAAGCGGCGTCCGCTAGC | PstⅠ |  |  |
| ORP2AS-C-F | ATGGATCCATGAATGAGAATCTCGTCAAAGAG | BamHⅠ | pPZP211-35S-mCherry | 35S::mCherry-ORP2AS |
| ORP2AS-C-R | ATGTCGACCTAAGCGGCGTCCGCTAGCT | SalⅠ |  |  |
| *orp2a-1*-T-F | CATCTTTCACGTGGATGTGTG |  |  | *orp2a-1* identification |
| *orp2a-1*-T-R | CTCCATCATCGAACTCCAGAG |  |  |  |
| PH crispr-F | GATTGGACTAATTTTGGTAAAGGA |  |  | CRISPR |
| PH crispr-R | AAACTCCTTTACCAAAATTAGTCC |  |  |  |
| ORD crispr-F | GATTGATCAAAGACAACGTTGGAA |  |  | CRISPR |
| ORD crispr-R | AAACTTCCAACGTTGTCTTTGATC |  |  |  |
| ACTIN2-Q-F | AGGCCAACAGAGAGAAGATG |  |  | Real-time PCR |
| ACTIN2-Q-R | CAGCACAATACCGGTTGTAC |  |  |  |
| F1 | GAACCAATATCATCCCTCC |  |  | *orp2a-1* Real-time PCR |
| R1 | CAGTGGAAGCATATCCAGAG |  |  |  |
| PH-Q-F | GGATAGTTGTAGTGGTCGC |  |  | Real-time PCR |
| PH-Q-R | GAAGGGTCTTTGTTGCTGTG |  |  |  |
| ORD-Q-F | ACAGGAGATACTTCCTCCC |  |  | Real-time PCR |
| ORD-Q-R | CGCTTTCACCTTGTCTCTC |  |  |  |
| 1-183aa-F | ATATTATGGCCATGGAGGCCATGCGGGTTAAAGAGTTACATCC | Sfi1 | AD | Y2H |
| 1-183aa-R | ATATTATGGCCTCCATGGCCTGACCGCAGAGGAAAGATAC | Sfi1 |  |  |
| 1-316aa-R | ATATTATGGCCTCCATGGCCTTCAGAACCAATGTTAGGTTCG | Sfi1 | AD | Y2H |
| 341-721aa-F | ATATTATGGCCATGGAGGCCATGGGTGTTAGTCTTTGGTCTATG | Sfi1 | AD | Y2H |
| 341-721aa-R | ATATTATGGCCTCCATGGCCAGCGGCGTCCGCTAGCTCTT | Sfi1 |  |  |
| 293-721aa-F | ATATTATGGCCATGGAGGCCATGGAGGAAGATGAACCTTCC | Sfi1 | AD | Y2H |
| ΔFF-F | CCTTCCACGACACAAAGGAGAACGAACCTAACATTGGTTC |  | AD | Y2H |
| ΔFF-R | GAACCAATGTTAGGTTCGTTCTCCTTTGTGTCGTGGAAGG |  |  |  |
| MutFF-F | CCACGACACAAAGGAGGCCGCTAACGAACCTAACATTG |  | AD | Y2H |
| MutFF-R | CAATGTTAGGTTCGTTAGCGGCCTCCTTTGTGTCGTGG |  |  |  |
| ORP2A-LUC-F | ATGGTACCATGCGGGTTAAAGAGTTACATC | Kpn1 | nLUC | LCI |
| ORP2A-LUC-R | ATGTCGACAGCGGCGTCCGCTAGCTCTT | Sal1 |  |  |
| VAP27-F | ATATTATGGCCATGGAGGCCATGAGTAACATCGATCTGATTG | Sfi1 | BD | Y2H |
| VAP27-R | ATATTATGGCCTCCATGGCCTTTGTCCTCTTCATAATGTATCCC | Sfi1 |  |  |
| VAP27-LUC-F | ATGGATCCATGAGTAACATCGATCTGATTG | BamHⅠ | cLUC  pGEX4T-1 | LCI  Pull-down |
| VAP27-LUC-R | ATGTCGACTTATGTCCTCTTCATAATGTATCC | Sal1 |  |  |
| RGS1-MY2H-F | ATATTATGGCCATTACGGCCATGGCGAGTGGATGTGCTC | Sfi1 | pPR3-suc | Y2H |
| RGS1-MY2H-R | ATATTATGGCCGAGGCGGCCTTACCGGGACTACTGCATCTG | Sfi1 |  |  |
| RGS1-FLUC-F | ATGGTACCATGGCGAGTGGATGTGCTC | Kpn1 | cLUC | LCI |
| RGS-LUC-R | ATGTCGACACCGGGACTACTGCATCTG | Sal1 |  |  |
